# GWAS of serum ALT and AST reveals an association of *SLC30A10* Thr95Ile with hypermanganesemia symptoms

**DOI:** 10.1101/2020.05.19.104570

**Authors:** Lucas D. Ward, Ho-Chou Tu, Chelsea B. Quenneville, Shira Tsour, Alexander O. Flynn-Carroll, Margaret M. Parker, Aimee M. Deaton, Patrick A. J. Haslett, Luca A. Lotta, Niek Verweij, Manuel A. R. Ferreira, Regeneron Genetics Center, Geisinger-Regeneron DiscovEHR Collaboration, Aris Baras, Gregory Hinkle, Paul Nioi

## Abstract

To investigate mechanisms of hepatocellular damage, we performed genome-wide association studies (GWAS) on alanine aminotransferase (ALT) and aspartate aminotransferase (AST) serum activities across 411,048 subjects from four ancestry groups in the UK Biobank, and found 100 loci associating with both enzymes. The rare missense variant *SLC30A10* Thr95Ile (rs188273166) associates with a larger elevation in ALT and AST than any other variant tested and this association also replicates in the DiscovEHR study. SLC30A10 excretes manganese from the liver to the bile duct, and rare homozygous loss of function causes the syndrome hypermanganesemia with dystonia-1 (HMNDYT1) which involves cirrhosis. Consistent with hematological symptoms of hypermanganesemia, *SLC30A10* Thr95Ile carriers have increased hematocrit and risk of iron deficiency anemia. Carriers also have increased risk of extrahepatic bile duct cancer. These associations suggest that genetic variation in *SLC30A10* adversely affects more individuals than patients with diagnosed HMNDYT1.

## Introduction

Liver disease remains an area of high unmet medical need, causing 3.5% of deaths worldwide, and the burden of liver disease is rising rapidly, driven mainly by increasing rates of nonalcoholic fatty liver disease (NAFLD)^1,2^. Better characterizing the genetic determinants of liver disease may lead to new therapies^3^. In addition, liver injury is a common side effect of drugs, and is a frequent reason that drugs fail to progress through the development pipeline; understanding the molecular mechanisms of liver injury can aid in preclinical drug evaluation to anticipate and avoid off-target effects^4,5^.

Circulating liver enzymes are sensitive biomarkers of liver injury; in particular, alanine aminotransferase (ALT) and aspartate aminotransferase (AST) are released into the circulation during damage to hepatocyte membranes^6,7^. The activities of circulating liver enzymes have been found to be highly heritable^6,8-12^, and variation even within the normal reference range is predictive of disease^13^. Accordingly, genome-wide association studies (GWAS) of activities of circulating liver enzymes across large population samples have proven powerful for understanding the molecular basis of liver disease^6,14-24^. Combined GWAS of ALT and AST have previously revealed genetic associations providing potential therapeutic targets for liver disease such as *PNPLA3*^25^ and *HSD17B13*^26^. To further study the genetics of hepatocellular damage, we performed GWAS on serum activities of ALT and AST in 411,048 subjects, meta-analyzed across four ancestry groups in the UK Biobank (UKBB).

## Results

### Discovery of ALT- and AST-associated loci by GWAS

We performed a GWAS of ALT and AST in four sub-populations in the UKBB (demographic properties, **Supplementary Table 1;** sample sizes, number of variants tested, and λ_GC_ values, **Supplementary Table 2;** genome-wide significant associations, **Supplementary Table 3**; Manhattan and QQ plots for each enzyme and sub-population, **Supplementary Figures 1 and 2**). After meta-analyzing across sub-populations to obtain a single set of genome-wide p-values for each enzyme (Manhattan plots, **Figure 1**), we found 244 and 277 independent loci associating at p < 5 × 10^−8^ with ALT and AST, respectively, defined by lead single nucleotide polymorphisms (SNPs) or indels separated by at least 500 kilobases and pairwise linkage disequilibrium (LD) r^2^ less than 0.2. Enzyme activities were strongly associated with coding variants in the genes encoding the enzymes, representing strong protein quantitative trait loci in *cis* (cis-pQTLs). For example rs147998249, a missense variant Val452Leu in *GPT* (glutamic-pyruvic transaminase) encoding ALT, strongly associates with ALT (p < 10^−300^) and rs11076256, a missense variant Gly188Ser in *GOT2* (glutamic-oxaloacetic transaminase 2) encoding the mitochondrial isoform of AST, strongly associates with AST (p = 6.3 × 10^−62^). While these strong *cis*-pQTL effects validated our ability to detect direct genetic influences on ALT and AST, the aim of this study was to detect genetic determinants of liver health that have downstream effects on both ALT and AST due to hepatocellular damage; therefore we focused the remainder of our analyses only on the variants associated with serum activity of both enzymes (labeled with black text on **Figure 1**).

**Figure 1:**
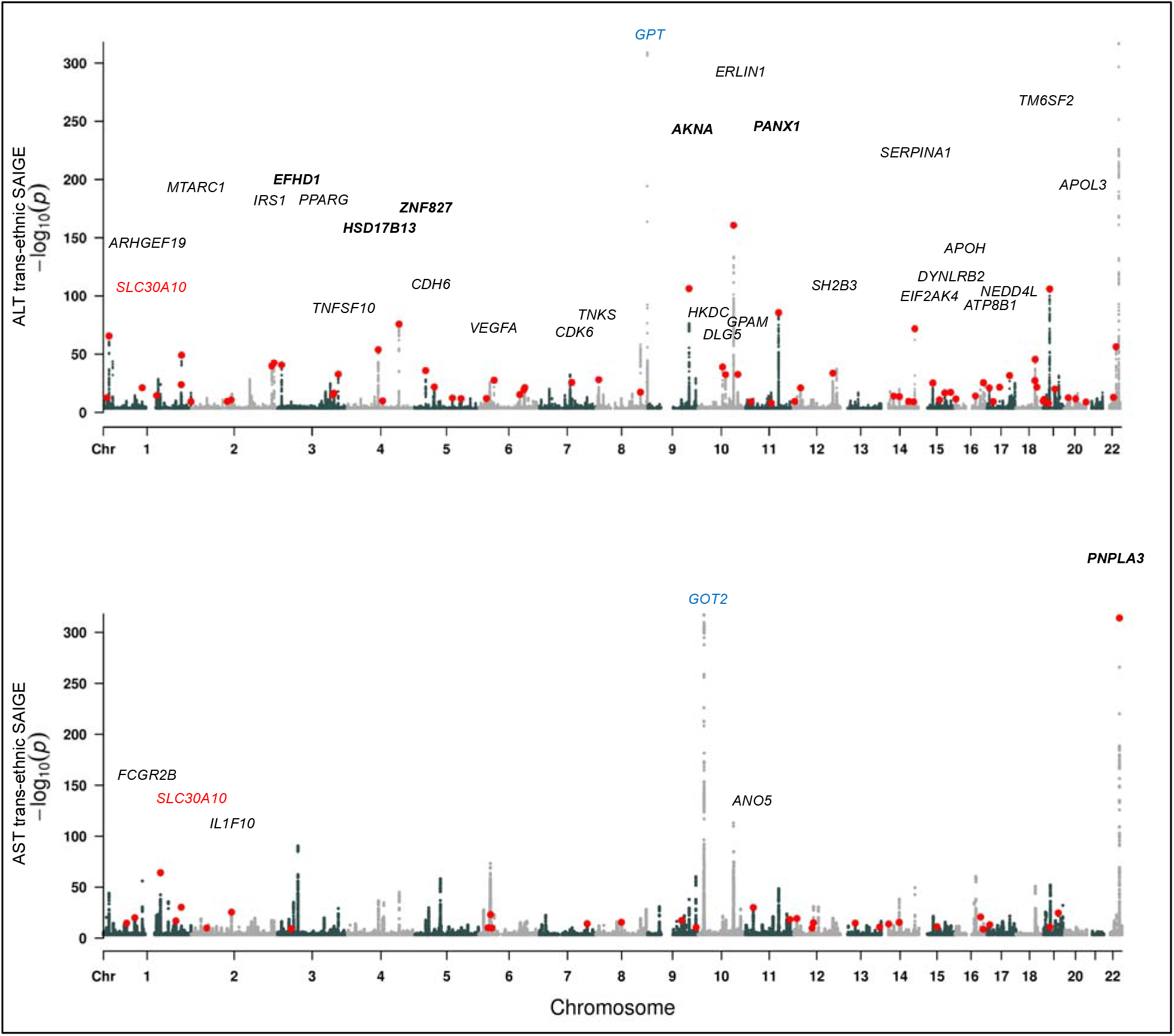
Manhattan plots showing trans-ancestry GWAS results for ALT and AST. Red dots indicate lead variants for shared signals between the two GWAS; for clarity, the shared signals are marked only once, on the plot for the GWAS in which the more significant association is detected. Cis-pQTLs (at *GPT* and *GOT2*) are labeled in blue. Loci with shared signals are labeled (for clarity, only when p < 10-25 and only on the GWAS for which the association is most significant). Loci previously reported to associate with both ALT and AST are named in bold. *SLC30A10*, the main topic of this report, is labeled in red on both plots.

Focusing only on loci with both ALT and AST GWAS signals (lead variants from either GWAS were identical or shared proxies with r^2^ ≥ 0.8), we found a total of 100 independent loci associated with both enzymes (**Figure 2, Supplementary Table 4**). As expected, effect sizes on ALT and AST at these loci were highly correlated (r = 0.98), and at all 100 loci the direction of effect on ALT and AST was concordant. Of these 100 loci, six were coincident or in strong LD with a published ALT or AST variant in the EBI-NHGRI GWAS Catalog, and 15 were within 500kb of a published ALT or AST variant; 33 of the loci harbored a missense or predicted protein-truncating variant; and of the remaining 67 entirely noncoding loci, 19 were coincident or in strong LD with the strongest eQTL for a gene in liver, muscle, or kidney suggesting that effects on gene expression may drive their associations with ALT and AST. A majority (70 of the 100 loci) were shared with a distinct published association in the GWAS Catalog, suggesting pleiotropy with other traits.

**Figure 2:**
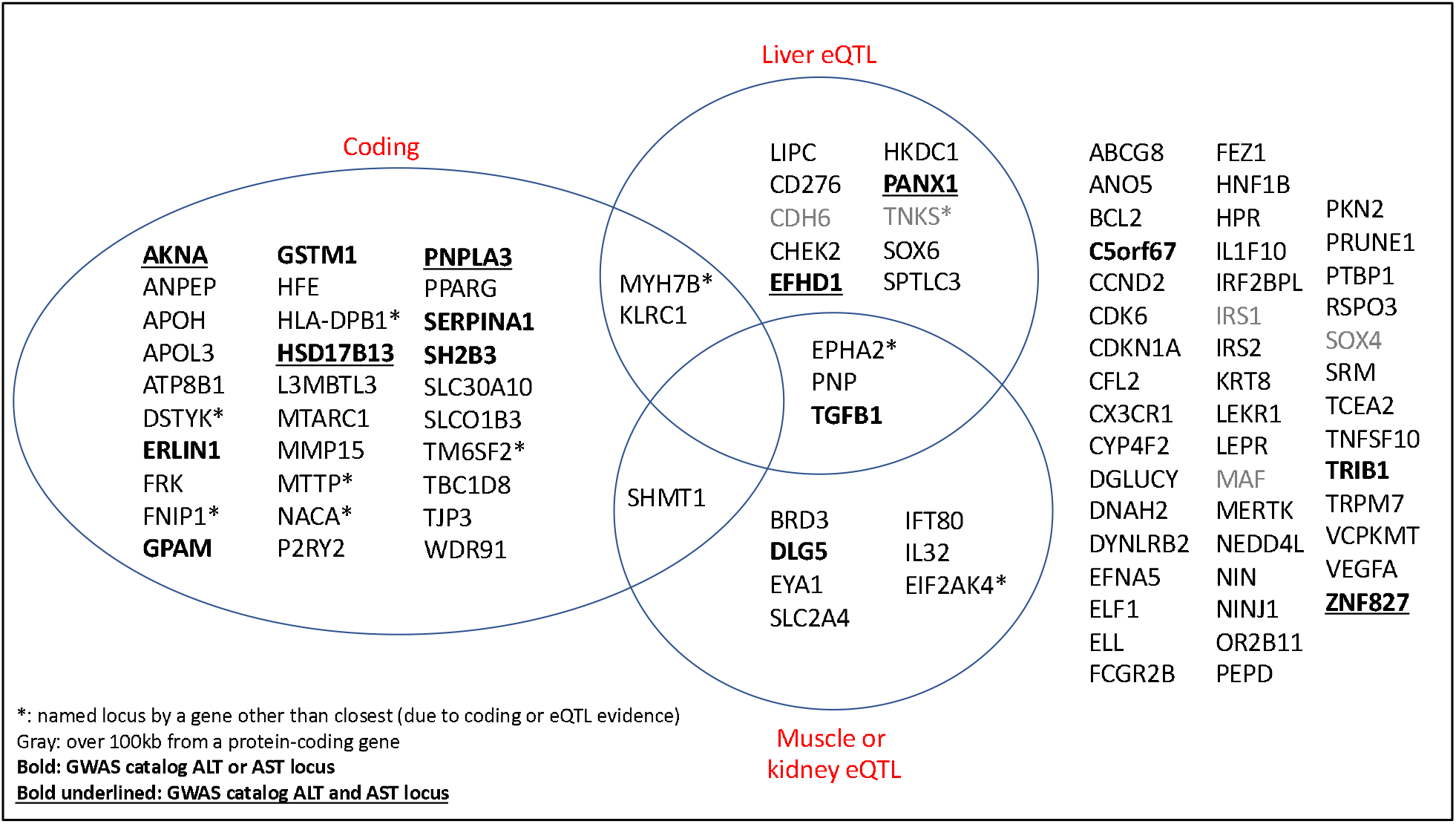
Classification of the top ALT- and AST-associated loci based on annotations. Unless otherwise noted with an asterisk, loci are named by the closest protein-coding gene. “Coding” indicates that one of the variants linked to the lead variant is predicted to have a moderate or high impact on a protein-coding gene. “Liver eQTL” and “Muscle or kidney eQTL” indicate that one of the variants linked to the lead variant is the strongest eQTL for a gene in those tissues by GTEx.

Comparing the effect sizes of all lead variants (**Supplementary Table 4**), the strongest estimated effect was a novel association: rs188273166, a rare (MAF in White British = 0.12%) missense variant (Thr95Ile) in *SLC30A10*, associated with a 4.2% increase in ALT (95% CI, 4.6% to 7.1%; p = 1.6 × 10^−24^) and 5.9% increase in AST (95% CI: 3.4% to 5.0%; p = 4.9 × 10^−31^). Because Thr95Ile is coding and not strongly linked to any other variants, we considered it likely to be the causal variant driving the association at the *SLC30A10* locus. *SLC30A10* encodes a manganese efflux transporter (solute carrier family 30 member 10, also known as zinc transporter 10 or ZnT10)^27,28^. Loss-of-function mutations in *SLC30A10* have been reported to cause a rare recessive syndrome, hypermanganesemia with dystonia 1 (HMNDYT1), characterized by cirrhosis, dystonia, parkinsonism, polycythemia, and hypermanganesemia^28-34^. The next strongest effect on either enzyme was rs28929474, a missense variant (the Pi-Z allele) in *SERPINA1* (serpin family A member 1) which causes alpha-1 antitrypsin deficiency (AATD) in its homozygous state^35^, associated with a 2.5% increase in ALT (95% CI, 2.1% to 2.8%; p = 1.4 × 10^−72^) and 1.3% increase in AST (95% CI, 1.1% to 1.5%; p = 3.2 × 10^−50^); AATD manifests with both lung and liver damage. The most statistically significant association with either enzyme was rs738409, a common missense variant (Ile148Met) in *PNPLA3* (patatin like phospholipase domain containing 3) known to strongly increase risk of liver disease^25^; it is associated with a 2.2% increase in ALT (95% CI, 2.1% to 2.3%; p < 10^−300^) and a 1.3% increase in AST (95% CI, 1.3% to 1.4%; p < 10^−300^).

We observed significant heterogeneity in effects between sexes for both enzymes (Cochran’s Q test p < 0.05/100 for both enzymes) for three of the lead variants: rs9663238 at *HKDC1* (stronger effects in women), rs28929474 at *SERPINA1* (stronger effects in men), and rs1890426 at *FRK* (stronger effects in men) (**Supplementary Table 5**).

We tested the 100 lead variants from the ALT and AST GWAS analysis for association with a broad liver disease phenotype (ICD10 codes K70-77; 14,143 cases and 416,066 controls), meta-analyzing liver disease association results across all four sub-populations (**Supplementary Table 4)**. Of the 100 lead variants, 28 variants associate with liver disease with p < 0.05. As expected, variants associated with an increase in ALT and AST tend to be associated with a proportional increase in liver disease risk (across all lead variants, Pearson correlation of betas r = 0.82 for both enzymes; **Figure 3**). Liver disease is found more frequently in our sample of carriers of *SLC30A10* Thr95Ile (rs188273166), proportional with the observation of increased ALT and AST (OR = 1.47); however, owing to the small sample size of carriers and liver disease cases, we are underpowered to confidently determine whether this high point estimate is due to chance (although the 95% CI from the PLINK analysis used to estimate effects does not include OR = 0, the p value from the SAIGE analysis which more accurately controls for Type I error in highly unbalanced case-control studies is 0.07; see Methods).

**Figure 3:**
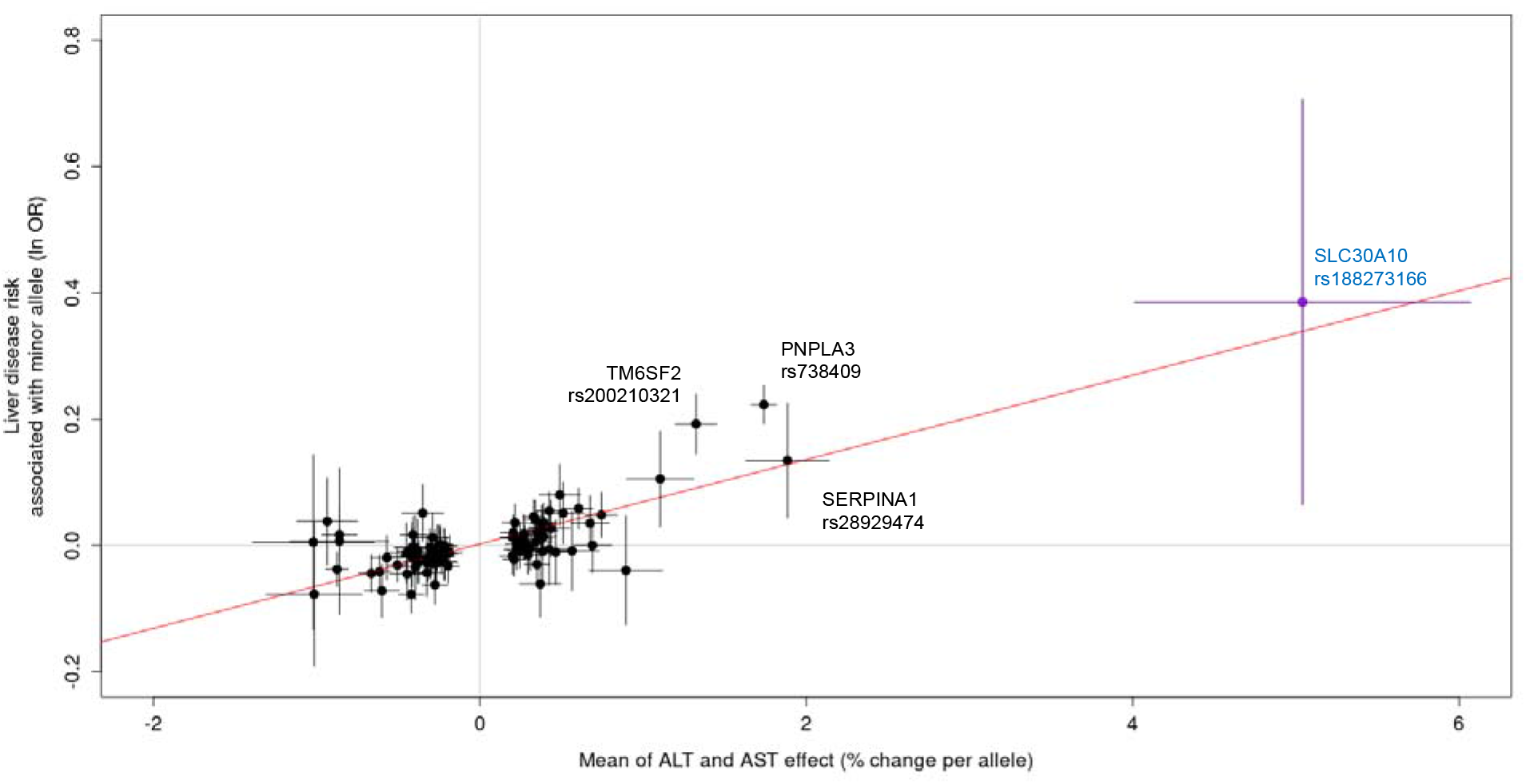
Comparison of effect of lead variants on ALT and AST with effect on liver disease. Effect sizes (beta from the regression of units of log_10_ ALT and AST, equivalent to percent change; and beta from the liver disease regression, equivalent to natural log of the odds ratio) are from PLINK analysis; error bars are 95% confidence intervals on these effect sizes from PLINK.

Because *SLC30A10* Thr95Ile had the strongest effect on ALT and AST of all of our lead variants and has not been reported as being associated with any phenotypes in the literature, we centered the following analyses on better understanding its function.

### Validation of *SLC30A10* Thr95Ile genotype

Because rare variants are especially prone to errors in array genotyping^36^, we sought to validate the array genotype calls for *SLC30A10* Thr95Ile in a subset of 301,473 individuals who had also been exome sequenced (**Supplementary Table 6**). The only individual homozygous for the minor (alternate) allele by array was confirmed by exome sequencing; no further homozygotes were identified. Of 702 individuals called as heterozygous for Thr95Ile by array data who had exome data available, 699 (99.6%) were confirmed heterozygous by exome sequencing, while three were called homozygous reference by exome sequencing, suggesting an error either in the array typing or exome sequencing for these three individuals. Overall, these results demonstrate high concordance between array and exome sequencing, implying highly reliable genotyping.

### Magnitude of ALT and AST elevation in *SLC30A10* Thr95Ile carriers

After establishing the association between *SLC30A10* Thr95Ile and ALT and AST, we sought to further explore the relationship between genotype and enzyme activity levels to understand clinical relevance. Carriers of Thr95Ile had a mean ALT of 27.37 U/L vs 23.54 U/L for noncarriers, and a mean AST of 28.85 U/L vs 26.22 U/L for noncarriers. Counting individuals with both ALT and AST elevated above 40 U/L, a commonly-used value for the upper limit of normal (ULN)^7^, 5.6% of carriers vs 3.6% of noncarriers had both enzymes elevated at the time of their UK Biobank sample collection, an increased relative risk of 58% (Fisher’s p = 8.1 × 10^−4^) (**Supplementary Table 7**).

### Drinking behavior in *SLC30A10* Thr95Ile carriers

The *SLC30A10* Thr95Ile has not been reported as associating with drinking behavior by any of the available studies in the GWAS Catalog. We used the drinking questionnaire taken by UK Biobank participants to assess drinking status at enrollment of *SLC30A10* Thr95Ile carriers (current, former, or never drinkers.) While the rate of current drinkers is higher among carriers vs non-carriers in the entire biobank (93.7% vs 91.7%, Fisher’s p = 0.019) (**Supplementary Table 7**), this association is highly confounded by genetic ancestry and country of birth (**Supplementary Table 8**). Limiting to the White British subpopulation and individuals born in England, the rate of current drinking is not detectably different among carriers (94.0% vs 93.4%, Fisher’s p = 0.57) while the rate of individuals with elevation of both ALT and AST over the ULN remains significant (5.5% vs. 3.5%, Fisher’s p = 4.6 × 10^−3^) (**Supplementary Table 7**).

### Replication of ALT and AST associations

The initial association of rs188273166 with ALT and AST was identified in the White British population. To replicate this association in independent cohorts, we first identified groups besides the White British subpopulation harboring the variant in the UKBB. The only two other populations with a substantial number of *SLC30A10* Thr95Ile carriers were individuals identifying as Other White and as White Irish (**Supplementary Table 8**); we tested for association with ALT and AST in these subpopulations. We then tested the association in two independent cohorts from the DiscovEHR collaboration between the Regeneron Genetics Center and the Geisinger Health System^37^. Meta-analyzing the association results across these four groups (N = 132,992 and N = 131,646, respectively) confirmed the Thr95Ile association with increased ALT and AST (p = 6.5 × 10^−5^ and p = 5.4 × 10^−6^, respectively) (**Supplementary Figure 3, Supplementary Table 9**). We also searched repositories of available complete summary statistics for ALT and AST GWAS and found two prior studies that reported associations^21,38^. Although these studies were underpowered to detect significant associations and were not reported in units that allowed their inclusion in our replication analysis, they were consistent with increases in both enzymes (**Supplementary Table 10**).

### Independence of *SLC30A10* Thr95Ile from neighboring ALT and AST associations

Because we applied distance and LD pruning to the results of the genome-wide scan to arrive at a set of lead variants, it was unclear how many independent association signals existed at the *SLC30A10* locus. Revisiting trans-ancestry association results in a window including 1 Mb flanking sequence upstream and downstream of *SLC30A10* revealed 76 variants with genome-wide significant associations with both ALT and AST (**Figure 4**). These 76 variants clustered into three loci: *SLC30A10* (only Thr95Ile, rs188273166); *MTARC1* (mitochondrial amidoxime reducing component 1, lead variant rs2642438 encoding missense Ala165Thr, previously reported to associate with liver disease and liver enzymes^39^, and six additional variants together spanning 68 kilobases); and *LYPLAL1-ZC3H11B* (intergenic region between lyophospholipase like 1 and zinc finger CCCH-type containing 11B, with array-genotyped variant rs6541227 and 67 imputed variants spanning 46 kilobases), a locus previously reported to associate with non-alcoholic fatty liver disease (NAFLD)^40^.

**Figure 4:**
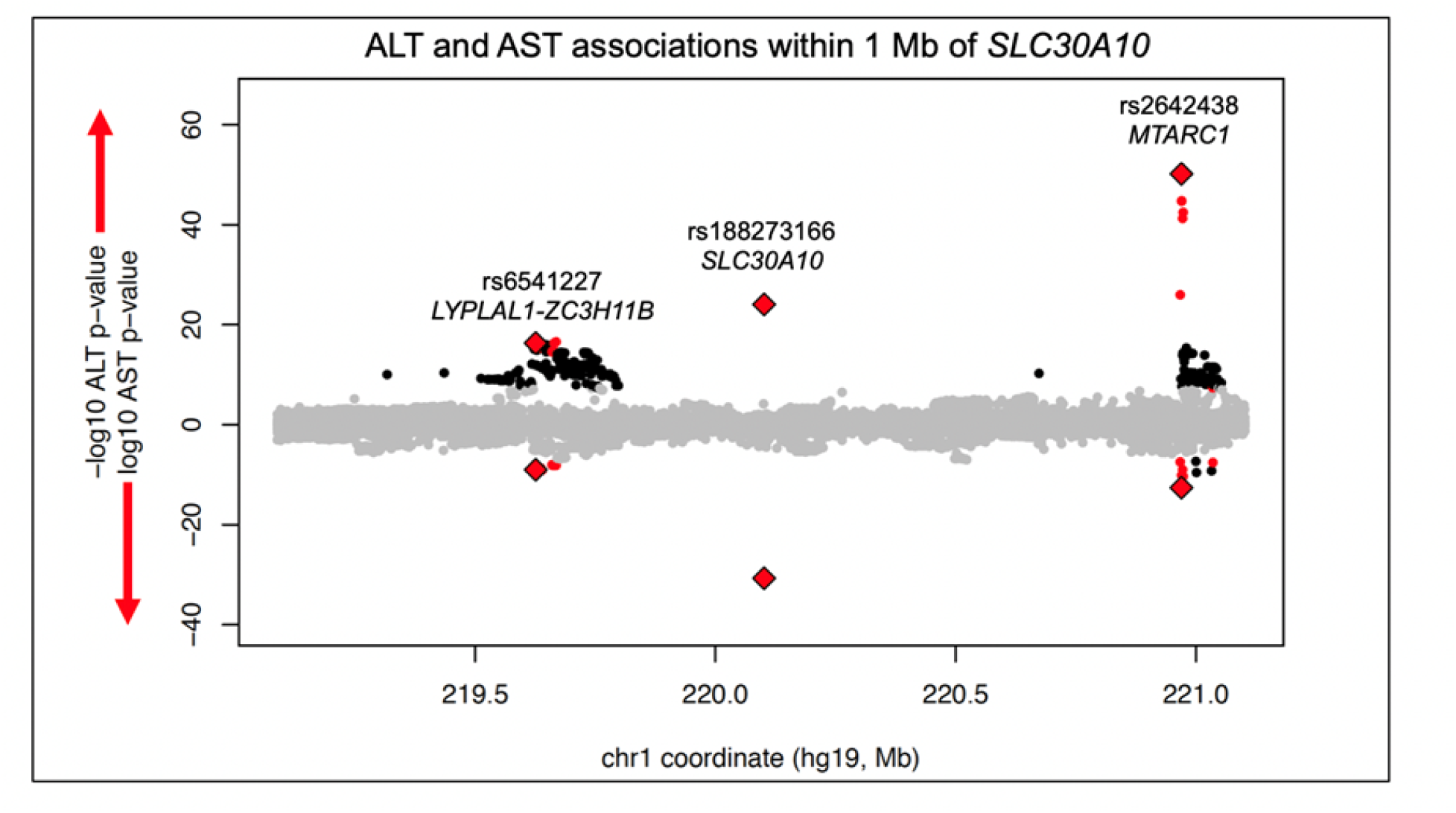
Miami plot of trans-ancestry SAIGE GWAS results within 1 Mb of the gene body of *SLC30A10*. ALT associations are shown in the positive direction and AST associations in the negative direction. Variant associations reaching genome-wide significance for one enzyme are colored black; for both enzymes, colored red; directly-genotyped variants significant for both enzymes, red diamonds.

To test for independence between these three loci, we performed ALT and AST association test for each of the three array-typed variants while including the genotype of either one or both of the others as covariates. Associations were similar in these conditional analyses, suggesting that each of these three associations are not confounded by linkage disequilibrium with the other regional association signals (**Supplementary Table 11**.) Therefore, the *SLC30A10* Thr195Ile association is statistically independent of the associations at neighboring loci. This statistical independence of the liver enzyme associations does not preclude a long-distance regulatory interaction between the three loci; for example, rs188273166, despite encoding an amino acid change in *SLC30A10*, could conceivably influence transcription of *MTARC1*, and rs6541227, despite being nearest to *LYPLAL1* and *ZC3H11B*, may influence transcription of *SLC30A10*. However, these three variants are not detected as liver eQTLs for the genes at neighboring loci in published data^41^.

### Linkage of Thr95Ile to GWAS variants at *SLC30A10*

A GWAS of circulating toxic metals^42^ discovered an association between a common intronic variant in *SLC30A10* (rs1776029; MAF in White British, 19.5%) and blood manganese levels, where the reference allele – which is the minor allele – is associated with increased circulating manganese. We calculated linkage disequilibrium statistics between rs1776029 and Thr95Ile and found that the minor allele of Thr95Ile (A) was in almost perfect linkage with the minor allele of rs1776029 (A) (r2 = 0.005, D’ = 0.98); Thr95Ile (rs188273166) is 154 times more frequent among carriers of at least one copy of the minor allele of common variant rs1776029 (95% CI = 84 – 325; Fisher’s p < 2.2 × 10^−16^). These results suggest that the previously reported association of rs1776029 with circulating manganese may be partially or completely explained by linkage with Thr95Ile (**Supplementary Table 12**); however, genotypes of Thr95Ile in the manganese GWAS or manganese measurements in the UK Biobank would be needed in order to perform conditional analysis or directly measure association of Thr95Ile with serum manganese. We then systematically tested nearby variants reported in the GWAS Catalog for any phenotype for linkage to Thr95Ile, measured by high |D’|. Combining GWAS Catalog information and |D’| calculations, we find nearly perfect linkage (|D’| > 0.90) between rs188273166-A (rare missense Thr95Ile) with rs1776029-A (intronic), rs2275707-C (3’UTR), and rs884127-G (intronic), all within the gene body of *SLC30A10* (**Supplementary Table 13**). In addition to increased blood Mn^42^, these three common alleles have been associated with decreased magnesium/calcium ratio in urine^43^, decreased mean corpuscular hemoglobin (MCH)^44-46^, increased red blood cell distribution width^44-46^, and increased heel bone mineral density (BMD)^46-49^. A recent study, not yet in the GWAS catalog, reported an association between another common intronic variant in *SLC30A10* (rs759359281; MAF in White British, 5.6%) and liver MRI-derived iron-corrected T1 measures (cT1)^50^. However, the reported cT1-increasing allele of rs759359281, which is the minor allele, is in complete linkage (D’ = 1) with the major allele of Thr95Ile (rs188273166); in other words, the cT1-increasing allele and Thr95Ile liver disease risk allele occur on different haplotypes, suggesting that the mechanism of this reported cT1 association is independent of Thr95Ile.

### Phenome-wide associations of *SLC30A10* Thr95Ile

To explore other phenotypes associated with *SLC30A10* Thr95Ile, we tested for association with 135 quantitative traits and 4,398 ICD10 diagnosis codes within the White British population (**Supplementary Tables 14 and 15**). We were particularly interested in testing associations with phenotypes related to HMNDYT1, the known syndrome caused by homozygous loss of function of SLC30A10. Besides ALT and AST elevation, rs188273166 was associated with other indicators of hepatobiliary damage such as decreased HDL cholesterol and apolipoprotein A (ApoA)^51^, decreased albumin, and increased gamma glutamyltransferase (GGT). Other phenome-wide significant quantitative trait associations were increases in hemoglobin concentration and hematocrit (**Table 1**); increased hematocrit, or polycythemia, is a known symptom of HMNDYT1. Liver iron-corrected T1 by MRI (cT1), although only measured in seven carriers, was above the population median value in all seven **(Supplementary Figure 4)**.

**Table 1:** Association results of *SLC30A10* Thr95Ile with selected quantitative traits. Association tests were performed in the White British population. P values are from SAIGE analysis; effect size estimates are from PLINK analysis. (***) indicates genome-wide significance p < 5 × 10^−8^; (**) indicates phenome-wide significance of p < 3.7 × 10^−4^; (*) indicates nominal significance of p < 0.05. All traits are rank-based inverse-normal transformed and effect size is in units of standard deviations of the transformed values.

| Quantitative trait | Association with *SLC30A10* Thr95Ile |  |  |  |  |  |
| --- | --- | --- | --- | --- | --- | --- |
|  | P | signif. | Effect (SD) | 95% CI lower | 95% CI upper | N |
| Aspartate aminotransferase | 2.90E-32 | *** | 0.38 | 0.31 | 0.44 | 387770 |
| Alanine aminotransferase | 1.95E-25 | *** | 0.32 | 0.25 | 0.38 | 389063 |
| HDL cholesterol | 1.18E-15 | *** | -0.22 | -0.28 | -0.15 | 356214 |
| Apolipoprotein A | 7.25E-11 | *** | -0.19 | -0.25 | -0.12 | 354247 |
| Albumin | 2.12E-08 | *** | -0.17 | -0.24 | -0.10 | 356374 |
| Hemoglobin concentration | 8.11E-06 | ** | 0.09 | 0.04 | 0.15 | 396065 |
| Hematocrit (percentage) | 1.33E-05 | ** | 0.10 | 0.04 | 0.15 | 396065 |
| Gamma glutamyltransferase | 9.86E-05 | ** | 0.12 | 0.05 | 0.18 | 388997 |
| Heel BMD | 2.05E-03 | * | 0.13 | 0.05 | 0.22 | 237053 |
| Liver MRI: iron corrected T1 (cT1) | 3.31E-03 | * | 1.22 | 0.48 | 1.96 | 2407 |

The only phenome-wide significant associations of diagnoses with *SLC30A10* Thr95Ile were C24.0, extrahepatic bile duct carcinoma, and C22.1, intrahepatic bile duct carcinoma. There are eight Thr95Ile carriers of each type of cancer, and six carriers with both types of cancer, for a total of ten carriers (1% of the 1,001 total carriers in the White British population) with bile duct carcinoma. Strikingly, over 5% of individuals with extrahepatic bile duct carcinoma (8 in 148) carry Thr95Ile. (**Table 2, Supplementary Table 15**).

**Table 2:** association results of *SLC30A10* Thr95Ile with selected ICD10 diagnosis codes. P values are from SAIGE analysis; effect size estimates are from PLINK analysis (exponentiated betas from logistic regression). (***) indicates genome-wide significance p < 5 × 10^−8^; (**) indicates phenome-wide significance of p < 1.1 × 10^−5^; (*) indicates nominal significance of p < 0.05. Case counts are all limited to the White British subpopulation (N = 408,160), the only subpopulation with a substantial number of carriers (N = 1,001).

| Diagnosis | Association with *SLC30A10* Thr95Ile |  |  |  |  |  |  |  |  |  |
| --- | --- | --- | --- | --- | --- | --- | --- | --- | --- | --- |
|  | P | sig. | N cases | Prevalence | N cases (carriers) | Prevalence (carriers) | N cases expected (carriers) | OR | 95% CI |  |
| C24.0: Malignant neoplasm of extrahepatic bile duct | 2.21E-09 | *** | 148 | 0.04% | 8 | 0.80% | 0.4 | 18.2 | 7.4 | 44.7 |
| C22.1: Intrahepatic bile duct carcinoma | 6.22E-06 | ** | 424 | 0.10% | 8 | 0.80% | 1.0 | 6.9 | 3.1 | 15.5 |
| K83.0: cholangitis | 1.64E-05 | * | 1297 | 0.32% | 13 | 1.30% | 3.2 | 3.4 | 1.8 | 6.6 |
| K83.1: obstruction of bile duct | 1.12E-04 | * | 1585 | 0.39% | 13 | 1.30% | 3.9 | 2.7 | 1.4 | 5.3 |
| D50.9: Iron deficiency anemia, unspecified | 9.86E-04 | * | 14687 | 3.60% | 55 | 5.49% | 36.0 | 1.7 | 1.3 | 2.3 |
| K81.0: Acute cholecystitis | 1.50E-03 | * | 2270 | 0.56% | 14 | 1.40% | 5.6 | 2.8 | 1.6 | 4.8 |
| D50: Iron deficiency anemia | 4.02E-03 | * | 20073 | 4.92% | 69 | 6.89% | 49.2 | 1.5 | 1.1 | 1.9 |

Among hematological manifestations of HMNDYT1, iron deficiency anemia was enriched among carriers (OR = 1.5, 95% CI, 1.1 to 1.9; p = 4.0 × 10^−3^). Searching for neurological manifestations similar to HMNDYT1, we find no association with Parkinson’s disease or dystonia but note that, as with liver diseases, we are powered to exclude only strong effects because of the small case number for these traits (**Supplementary Table 15**).

The top non-cancer hepatobiliary associations with *SLC30A10* Thr95Ile were with K83.0, cholangitis; K83.1, obstruction of bile duct; and K81.0, acute cholecystitis. Because biliary diseases are risk factors for cholangiocarcinoma and co-occur with them in our data, we tested whether *SLC30A10* Thr95Ile was still associated with these biliary diseases, and the other selected quantitative traits and diagnoses, after removing the 148 individuals with extrahepatic bile duct cancer (**Supplementary Table 16, Supplementary Table 17**). All of the associations remained significant except for intrahepatic bile duct carcinoma.

To test whether the association with extrahepatic bile duct cancer was driven by a nearby association, and to assess other risk variants and the potential for false positives given the extreme case-control imbalance, we performed a GWAS of the phenotype; remarkably, *SLC30A10* Thr95Ile was the strongest association genomewide, with minimal evidence for systematic inflation of p-values (λ_GC_ = 1.05; **Figure 5, Supplementary Figure 5**).

**Figure 5.**
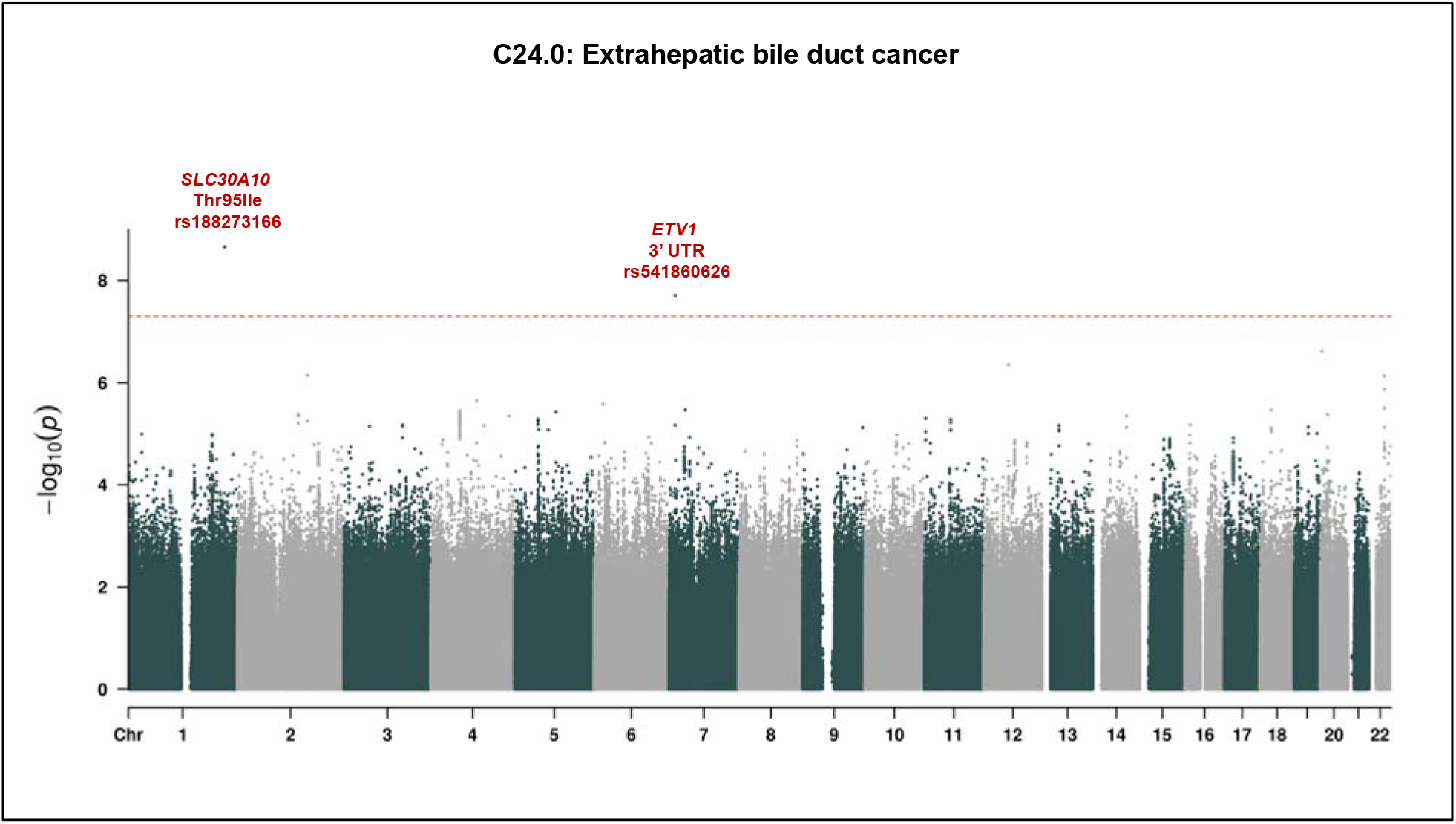
GWAS of extrahepatic bile duct cancer. P values are from SAIGE in White British. Dashed line is genome-wide significance level of p = 5 × 10^−8^.

### Replication of hematocrit association

A key result from the phenome-wide scan that was not related to hepatocellular damage was the association between *SLC30A10* Thr95Ile and increased hematocrit. Polycythemia is a symptom of HMNDYT1 mechanistically related to manganese overload. We meta-analyzed hematocrit values from the Other White and White Irish populations, the DiscovEHR data, and a non-UKBB population (INTERVAL Study) from a published meta-analysis of hematocrit values^45^, and found that the association replicated (N = 179,689, p = 0.013; **Supplementary Table 18**).

### *SLC30A10* expression in liver cell subtypes

Across organs, SLC30A10 is transcribed at the highest level in liver according to data from the GTEx Project^52^. The association of Thr95Ile with bile duct cancer led us to query expression of SLC30A10 in specific cell types within the liver using data from three single-cell RNA sequencing studies of liver^53-55^. These data show very low expression of SLC30A10 message in individual cells, but all studies detect expression in both hepatocytes and cholangiocytes (**Supplementary Table 19, Supplementary Figure 7**). Immunohistochemistry has established that SLC30A10 protein is present in hepatocytes and bile duct epithelial cells and localizes to the cholangiocyte plasma membrane, facing the lumen of the bile duct^32^.

### Bioinformatic characterization of *SLC30A10* Thr95Ile

To understand potential functional mechanisms of the Thr95Ile variant, we examined bioinformatic annotations of SLC30A10 Thr95Ile from a variety of databases. The UNIPROT database shows that Thr95Ile occurs in the third of six transmembrane domains and shares a domain with a variant known to cause HMNDYT1 (**Supplementary Figure 8**). Several *in silico* algorithms predict that Thr95Ile is a damaging mutation. The CADD (Combined Annotation Dependent Depletion) algorithm, which combines a broad range of functional annotations, gives the variant a score of 23.9, placing it in the top 1% of deleteriousness scores for genome wide potential variants. The algorithm SIFT, which uses sequence homology and physical properties of amino acids, predicts Thr95Ile as deleterious. The algorithm PolyPhen-2 gives Thr95Ile a HumDiv score of 0.996 (probably damaging), based on patterns of sequence divergence from close mammalian homologs, and a HumVar score of 0.900 (possibly damaging), based on similarity to known Mendelian mutations. Cross-species protein sequence alignment in PolyPhen-2 shows only threonine or serine at position 95 across animals. These properties suggest that Thr95Ile substitution ought to have an effect the function of the SLC30A10 protein.

### Characterization of *SLC30A10* variants *in vitro*

To test the protein localization of SLC30A10 harboring Thr95Ile as well as other variants, we created constructs with Thr95Ile (rs188273166) and the HMNDYT1-causing variants Leu89Pro (rs281860284) and del105-107 (rs281860285) and transfected these constructs into HeLa cells. Immunofluorescence staining revealed membrane localization for wild-type (WT) SLC30A10 which was abolished by the two HMNDYT1 variants, consistent with previous reports which showed that the HMNDYT1 variant proteins are mislocalized in the endoplasmic reticulum (ER)^56^. In contrast, Thr95Ile showed membrane localization similar to WT, suggesting that Thr95Ile does not cause a deficit in protein trafficking to the membrane (**Figure 6**).

**Figure 6:**
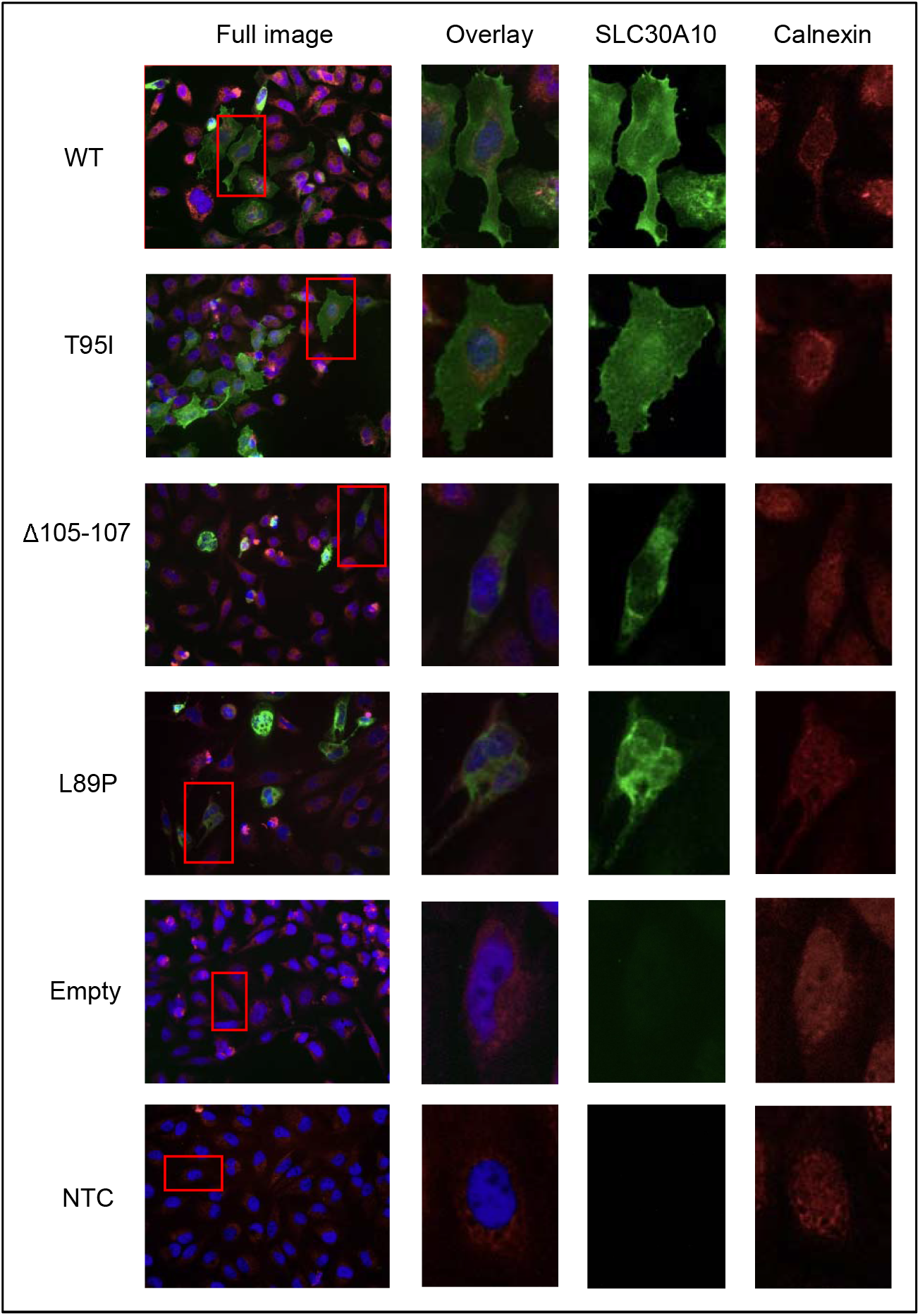
Immunofluorescence imaging of SLC30A10 protein constructs expressed in cultured HeLa cells. WT = wild type; T95I = Thr95Ile; Δ105-107 and L89P, HMNDYT1-causing variants reported previously [^56^]; empty = transfected with empty vector; NTC = non transfected control. Calnexin staining (in red) indicates the endoplasmic reticulum (ER).

## Discussion

### Expanded genetic landscape of risk for hepatocellular damage

Our trans-ancestry GWAS of ALT and AST reveals a broad genetic landscape of loci that modulate risk of hepatocellular damage or other diseases that cause increases in circulating ALT and AST, bringing the number of loci known to associate with serum activities of both enzymes from 10 (currently in the GWAS Catalog) to 100. Two loci had been previously reported in majority-European ancestry GWAS of ALT and AST as associating with both enzymes: *PNPLA3*^14,15,18-20,57-59^ and *HSD17B13*^14,18,26^; we detect ALT and AST signals at both of these loci. Broadening beyond majority-European ancestry GWAS, an additional eight loci have been previously identified: *PANX1*^18^, *ALDH2*^17,19^, *CYP2A6*^18,57^, *ABCB11*^18^, *ZNF827*^18^, *EFHD1*^18^, *AGER-NOTCH4*^18,60^, and *AKNA*^18^; we replicate four of these in our trans-ancestry GWAS (*PANX1, ZNF827, EFHD1*, and *AKNA*.) We are limited by the lack of diversity in the UK Biobank and expect that studies in more diverse populations will result in the discovery of new loci and alleles.

Among the novel loci are many that had been previously identified as risk loci for liver disease, but had never been explicitly associated through GWAS of both ALT and AST, such as *SERPINA1* (associated with alpha-1 antitrypsin deficiency^35^), *HFE* (homeostatic iron regulator, associated with hemochromatosis^61^), and *TM6SF2* (transmembrane 6 superfamily member 2, associated with NAFLD^62-64^). The *MTARC1* lead variant was discovered in a GWAS of cirrhosis, and then found to associate with lower ALT and AST^39^. Others are known to associate with risk of gallstones (*ABCG8, ANPEP*, and *HNF1B*)^65,66^ or increased GGT (*EPHA2, CDH6, DLG5, CD276, DYNLRB2*, and *NEDD4L*)^14,18^. Consistent with the fact that ALT and AST elevation can be caused by kidney or muscle damage, we detect an association with *ANO5* (anoctamin 5), which has been implicated in several autosomal recessive muscular dystrophy syndromes^67,68^, and several loci associated with expression of genes in muscle or kidney but not liver (*SHMT1, BRD3, DLG5, EYA1, IFT80, IL32, EIF2AK4*, and *SLC2A4*). We expect only a subset of the loci from this screen to be directly causally implicated in hepatocellular damage; many may predispose to a condition where liver damage is secondary or affect kidney or muscle, an important limitation of this approach.

The significant sex heterogeneity we observe at the ALT- and AST-associated loci *HKDC1, SERPINA1*, and *FRK* warrants further investigation and is consistent with a prior study that found significant genotype-by-sex interactions in the genetic architecture of circulating liver enzymes^6^. *HKDC1* has been associated with glucose metabolism and notably, this effect is specific to pregnancy^69,70^.

*SLC30A10* had not previously been identified in prior GWAS of circulating liver enzymes. Because *SLC30A10* Th95Ile is so rare, it is not surprising that these scans were underpowered to detect its large effect, due either to insufficient study size, lack of inclusion on the genotyping arrays used, or lack of power to impute its genotype. For example, in the UK Household Longitudinal Study (N = 5,458 and N = 5,321, respectively) effects were reported that were consistent with a strong effect size but were not statistically significant (**Supplementary Table 10**).

### Properties of *SLC30A10* Thr95Ile

The variant with the strongest predicted effect on ALT and AST, *SLC30A10* Thr95Ile (rs188273166), is a rare variant carried by 1,117 of the 487,327 array-genotyped participants in the UK Biobank. While Thr95Ile is found in some individuals of non-European ancestry, it is at much higher frequency in European-ancestry populations, with carrier frequency in our sample by UK country of birth ranging from a minimum of 1 in 479 people born in Wales to a maximum of 1 in 276 people born in Scotland (**Supplementary Table 5**). The increased frequency we see in European-ancestry populations is not merely due to those populations’ overrepresentation in the UK Biobank, but is also consistent with global allele frequency data catalogued in dbSNP^71^.

The Thr95Ile variant occurs in the third of six transmembrane domains of the SLC30A10 protein^72^, the same domain affected by a previously reported loss-of-function variant causing HMNDY1 (hypermanganesemia with dystonia 1), Leu89Pro (rs281860284)^56^ (**Figure 6**). In vitro, Leu89Pro abolishes trafficking of SLC30A10 to the membrane^56^, and another study pointed to a functional role of polar or charged residues in the transmembrane domains of SLC30A10 for manganese transport function^73^. Bioinformatic analysis suggests that Thr95Ile should impact protein function.

Our site-directed mutagenesis experiment of SLC30A10 shows that Thr95Ile, unlike reported HMNDYT1-causing variants, results in a protein that is properly trafficked to the cell membrane. Further biochemical studies will be required to investigate whether the Thr95Ile variant of SLC30A10 has reduced manganese efflux activity, or otherwise affects SLC30A10 stability, translation, or transcription.

### Comparison of *SLC30A10* Thr95Ile phenotypes to HMNDYT1 phenotypes

*SLC30A10* (also known as *ZNT10*, and initially identified through sequence homology to zinc transporters^27^) encodes a cation diffusion facilitator expressed in hepatocytes, the bile duct epithelium, enterocytes, and neurons^32^ that is essential for excretion of manganese from the liver into the bile and intestine^28,32^. Homozygous loss-of-function of *SLC30A10* was recently identified as the cause of the rare disease HMNDYT1, which in addition to hypermanganesemia and dystonia is characterized by liver cirrhosis, polycythemia, and Mn deposition in the brain^29-34,56^. Other hallmarks include iron depletion and hyperbilirubinemia. Mendelian disorders of *SLC30A10* and the other hepatic Mn transporter genes *SLC39A8* (solute carrier family 39 member 8, causing congenital disorder of glycosylation type IIn)^74^ and *SLC39A14* (solute carrier family 39 member 14, implicated in hypermanganesemia with dystonia 2)^75^, along with experiments in transgenic mice ^76,77^, have confirmed the critical role of each of these genes in maintaining whole-body manganese homeostasis^78^. Notably, while all three of the genes have Mendelian syndromes with neurological manifestations, only *SLC30A10* deficiency (HMNDYT1) is known to be associated with liver disease^78^.

We detect two key aspects of HMNDYT1 – increased circulating liver enzymes and increased hematocrit – exceeding phenome-wide significance in heterozygous carriers of *SLC30A10* Thre95Ile. Among other hepatic phenotypes that have been reported in HMNDYT1 cases, we also detect an association with anemia, but no evidence of hyperbilirubinemia. The neurological aspect of HMNDYT1, parkinsonism and dystonia, is not detectably enriched among Thr95Ile carriers; however, we have limited power and cannot exclude an enrichment. It is therefore intriguing to consider that carrier status of Thr95Ile may represent a very mild manifestation of HMNDYT1.

The quantitative trait with the largest effect associated with *SLC30A10* Thre95Ile is liver MRI cT1 (+1.2 SD; 95% CI, +0.5 to +2.0; p = 0.0032). Liver MRI cT1 has been recently explored as a non-invasive diagnostic of steatohepatitis and fibrosis^79,80^. However, MRI T1 signal has also been used to detect manganese deposition in the brain, and it is unclear the extent to which hepatic manganese overload could confound the association of liver cT1 with liver damage^81^.

### Comparison of Thr95Ile phenotypes to *SLC30A10* common variant phenotypes

Apart from rare variants in *SLC30A10* causing HMNDYT1, Thr95Ile can also be compared to common variants in *SLC30A10* that have been associated with phenotypes by GWAS (**Figure 7**). We find that the minor allele of Thr95Ile is in almost complete linkage with a common intronic variant associated with increased blood manganese. Other GWAS variants in almost perfect linkage with Thr95Ile associate with decreased MCH, increased RBC distribution width, decreased magnesium/calcium ratio, and increased heel bone mineral density (BMD). Decreased MCH could reflect the anemia experienced by HMNDYT1 patients, caused by the closely linked homeostatic regulation of manganese and iron^28^. Increased BMD may reflect the protective role of manganese in bone maintenance^82,83^. Looking for the subset of these phenotypes available in our scan of Thr95Ile, we do find a nominally significant increase in BMD but no detectable increase in MCH or erythrocyte distribution width. By contrast, we find that a common intronic variant in *SLC30A10* recently reported to associate with liver MRI cT1^50^ is in complete linkage with the major allele of Thr95Ile, suggesting an independent genetic mechanism but also providing independent evidence of the role of *SLC30A10* variants in liver health and/or hepatic manganese content.

**Figure 7:**
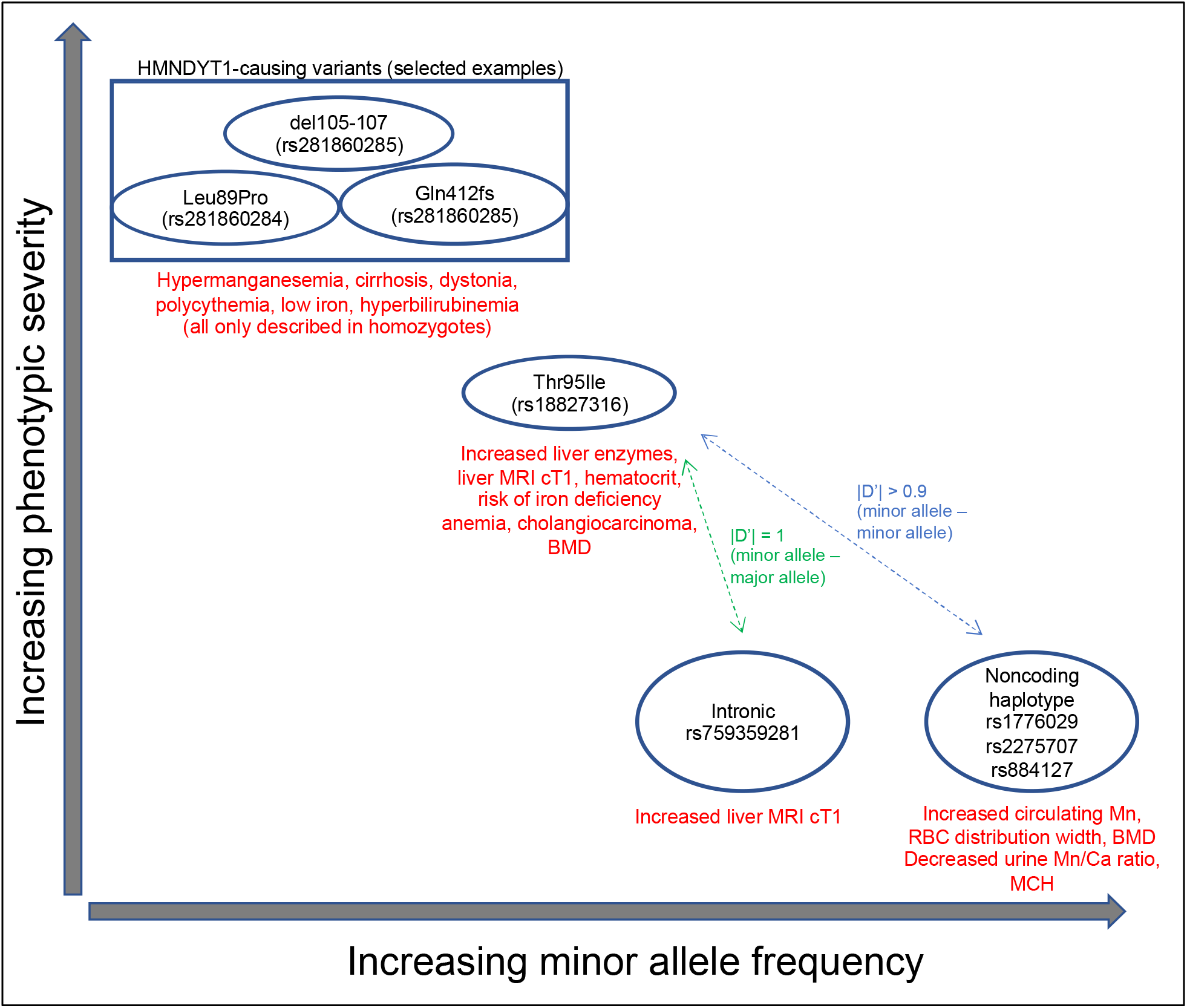
Relationship of Mendelian and common-variant GWAS phenotypes at the *SLC30A10* locus. Phenotypes are summarized for HMNDYT1-causing variants, *SLC30A10* Thr95Ile, and GWAS variants.

The linked GWAS variants may be interpreted through two mechanistic hypotheses: first, the associations may all be causally driven by Thr95Ile carriers in the studies, which the GWAS variants tag; alternatively, the associations may be driven by effects of the common variants themselves, which are noncoding but may influence *SLC30A10* (or another gene in cis) by modulating expression or post-transcriptional regulation; or some combination of both. To distinguish between these, measurements of Mn would need to be available to perform conditional analyses. If the GWAS variants have an effect independent of Thr95Ile, *SLC30A10* still seems likely (although not certain) to be the causal gene at the locus, due to the similarity in phenotypes to HMNDYT1 and Thr95Ile. A putative regulatory mechanism could be through transcriptional or post-transcriptional regulatory elements, as the haplotype includes a variant (rs2275707) overlapping both the 3’-UTR of *SLC30A10* and H3K4me1 histone modifications characteristic of enhancers active only in brain and liver^84^.

### Clinical relevance: manganese homeostasis in health and disease

Manganese (Mn) is a trace element required in the diet for normal development and function, serving as a cofactor and regulator for many enzymes. However, it can be toxic in high doses; because Mn(II) and Mn(III) can mimic other cations such as iron, it can interfere with systemic iron homeostasis and disrupt in other biochemical processes^85,86^; at the cellular level, it is cytotoxic and poisons the mitochondria by interfering with the electron transport chain enzymes that rely on Fe-S clusters^87^. The hallmark of occupational exposure through inhalation is neurotoxicity manifesting as parkinsonism and dystonia (manganism, or Mn intoxication)^85,86^. Neurotoxicity is an aspect of the Mendelian syndromes caused by loss of function of all three of the hepatic manganese transporters; interestingly, GWAS has also identified a common missense variant in *SLC39A8* as a risk factor for schizophrenia and many other diseases^88,89^; altered function of glycosyltransferases due to manganese overload in the neurons is a proposed mechanism for neurological manifestations of this variant^90^. Because manganese is excreted through the liver into the bile, increased circulating manganese secondary to liver damage may be a contributing factor to the neurological manifestations of chronic acquired hepatocerebral degeneration (CAHD)^91-93^. However, liver toxicity is not a hallmark of environmental or occupational exposure. Importantly, of the Mendelian syndromes of genes encoding manganese transporters, only *SLC30A10* (causing HMNDYT1) involves hepatic symptoms^78,94^. Hepatotoxicity in HMNDYT1 is thought to be due to cytotoxic manganese overload within hepatocytes; polycythemia is thought to be caused by upregulation of erythropoietin by manganese; and iron anemia through systemic dysregulation of iron homeostasis by excess manganese^94,95^. Our results suggest that polymorphism in *SLC30A10* is a risk factor for manganese-induced hepatocellular damage, polycythemia, and iron anemia in a much broader population beyond the rare recessive syndrome HMNDYT1.

The association of *SLC30A10* Thr95Ile with extrahepatic bile duct cancer was unexpected, as this disease has not been described in conjunction with HMNDYT1. Bile duct cancer (cholangiocarcinoma) is a rare disease (age-adjusted incidence of 1 – 3 per 100,000 per year); cirrhosis, viral hepatitis, primary sclerosing cholangitis, and parasitic fluke infection have been identified as risk factors^96,97^. It is unclear whether low levels of manganese in the bile, or high levels of manganese in the hepatocytes and bile duct epithelial cholangiocytes, could be directly carcinogenic; manganese-dependent superoxide dismutase (MnSOD, or SOD2) is a tumor suppressor^98^. A simpler possibility is that cytotoxic manganese overload in hepatocytes and cholangiocytes causes localized inflammation that predisposes to cancer through similar mechanisms as other hepatobiliary risk factors. We do detect an association with cholangitis, but the effect of this association is weaker than the association with cholangiocarcinoma. To our knowledge, *SLC30A10* Thr95Ile would be the strongest genetic cholangiocarcinoma risk factor identified to date, being carried by 5% of the extrahepatic bile duct cancer cases in the White British subset of the biobank. Because both *SLC30A10* Thr95Ile and extrahepatic bile duct cancer are exceedingly rare, validation of this association in either another very large biobank or in a cohort of cholangiocarcinoma patients will be necessary.

### Clinical relevance: genome interpretation

Currently, *SLC30A10* Thr95Ile (rs188273166) is listed as a variant of uncertain significance in the ClinVar database^99^. While the appropriate clinical management of carriers of *SLC30A10* Thr95Ile is unclear and would require further studies to determine whether monitoring of hepatobiliary function is warranted, evidence from HMNDYT1 patients has demonstrated that chelation therapy combined with iron supplementation is effective at reversing the symptoms of SLC30A10 insufficiency^100^. Further studies will be needed to define whether other damaging missense variants or protein-truncating variants in *SLC30A10*, including the variants known to cause HMNDYT1, also predispose to liver disease in their heterozygous state. Because we only observe one homozygous carrier of *SLC30A10* Thr95Ile in our data, further study will also be needed to understand the inheritance model of this association; we cannot determine whether risk in homozygotes is stronger than risk in heterozygotes, unlike cases of HMNDYT1 where identified cases have all experienced homozygous loss-of-function mutation.

More broadly, the case of *SLC30A10* fits a pattern of discoveries of recent discoveries showing that recessive Mendelian disease symptoms can manifest in heterozygous carriers of deleterious variants, blurring the distinction between recessive and dominant disease genes and bridging the gap between common and rare disease genetics^101,102^. These discoveries are possible only by combining massive, biobank-scale genotype and phenotype datasets such as the UK Biobank.

## Methods

### Sub-population definition and PC calculation

Sub-populations for analysis were obtained through a combination of self-reported ethnicity and genetic principal components. First, the White British population was defined using the categorization performed previously by the UK Biobank^103^ (Field 22006 value “Caucasian”); briefly, this analysis selected the individuals who identify as White British (Field 21000), performed a series of subject-level QC steps (to remove subjects with high heterozygosity or missing rate over 5%, removing subjects with genetic and self-reported sex discrepancies and putative sex chromosome aneuploidies, and removing subjects with second or first degree relatives and an excess of third degree relatives), performed Bayesian outlier detection using the R package aberrant ^104^ to remove ancestry outliers using principal components (PCs) 1+2, 3+4, and 5+6 (calculated from the global UK Biobank PCs stored in Field 22009), selected a subset of variants in preparation for PCA by limiting to directly-genotyped (info = 1), missingness across individuals < 2%, MAF > 1%, regions of known long range LD, and pruning to independent markers with pairwise LD < 0.1. Based on this procedure used by the UK Biobank to define the “White British” subset, we defined three additional populations, using other self-reported ancestry groups as starting points (Field 21000 values “Asian or Asian British”, “Black or Black British”, and “Chinese”). Principal components were estimated in PLINK using the unrelated subjects in each subgroup. We then projected all subjects onto the PCs. For the majority of downstream analyses (calculation of per-variant allele frequency and missingness thresholds, calculation of LD, and for association analyses performed in PLINK), just the unrelated subset of people in the subpopulation was used, the unrelated sets were used. The exception was association analyses performed in SAIGE^105^, a generalized mixed model method that allows inclusion of related individuals; for SAIGE, related individuals were retained in the subpopulations.

For validation in an independent subpopulation of the UK Biobank, two other self-reported ethnicity groups with a sufficient number of *SLC30A10* Thr95Ile carriers were assembled, who were not included in “White British” (Field 21000 values “White” subgroup “Irish”, and “White” subgroup “Any other white background” or no reported subgroup).

### Array genotype data for association analysis

Data were obtained from the UK Biobank through application 26041. Genotypes were obtained through array typing and imputation as described previously^103^. For genome-wide association analysis, variants were filtered so that imputation quality score (INFO) was greater than 0.8. Genotype missingness, Hardy-Weinberg equilibrium (HWE), and minor allele frequency (MAF) were then each calculated across the unrelated subset of individuals in each of the four sub-populations. For each sub-population a set of variants for GWAS was then defined by filtering missingness across individuals less than 2%, HWE p-value > 10^−12^, and MAF > 0.1%.

### Phenotype data

For genome-wide analysis, blood biochemistry values were obtained for ALT (Field 30620) and AST (Field 30650) and log_10_ transformed, consistent with previous genetic studies^14,106^, resulting in an approximately normal distribution.

For phenome-wide analysis, ICD10 codes were obtained from inpatient hospital diagnoses (Field 41270), causes of death (Field 40001 and 40002), the cancer registry (Field 40006), and general practitioner (GP) clinical event records (Field 42040). A selection of 135 quantitative traits was obtained from other fields (**Supplementary Table 14**), encompassing anthropomorphic measurements, blood and urine biochemistry, smoking, exercise, sleep behavior, and liver MRI; all were inverse rank normalized using the RNOmni R package^107^. All quantitative traits and cancer registry diagnoses were downloaded from the UK Biobank Data Showcase on March 17, 2020. The GP clinical events, inpatient diagnoses, and death registry were available in more detail or in more recent updates than was available through the Data Showcase and were downloaded as separate tables; data for GP clinical records were downloaded on March 30, 2019, data from the death registry was downloaded on June 12, 2020, and data from hospital diagnoses was downloaded on July 15, 2020.

### Genome-wide association studies of ALT and AST

Because of the high level of relatedness in the UK Biobank participants^108^, to maximize power by retaining related individuals we used SAIGE software package^105^ to perform generalized mixed model analysis for GWAS. A genetic relatedness matrix (GRM) was calculated for each sub-population with a set of 100,000 LD-pruned variants selected from across the allele frequency spectrum. SAIGE was run on the filtered imputed variant set in each sub-population using the following covariates: age at recruitment, sex, BMI, and the first 12 principal components of genetic ancestry (learned within each sub-population as described above). Manhattan plots and Q-Q plots were created using the qqman R package^109^. The association results for each enzyme were meta-analyzed across the four populations using the METAL software package^110^ using the default approach (using p-value and direction of effect weighted according to sample size.) To report p-value results, the default approach was used. To report effect sizes and standard errors, because the authors of the SAIGE method advise that parameter estimation may be poor especially for rare variants^111^, the PLINK software package v1.90^112^ was run on lead variants on the unrelated subsets of each subpopulation, and then the classical approach (using effect size estimates and standard errors) was used in METAL to meta-analyze the resulting betas and standard errors. All PLINK and SAIGE association tests were performed using the REVEAL/SciDB translational analytics platform from Paradigm4.

### Identifying independent, linked association signals between the two GWAS

Meta-analysis results for each enzyme were LD clumped using the PLINK software package, v1.90^112^ with an r^2^ threshold of 0.2 and a distance limit of 10 megabases, to group the results into approximately independent signals. LD calculations were made using genotypes of the White British sub-population because of their predominance in the overall sample. Lead variants (the variants with the most significant p-values) from these “r^2^ > 0.2 LD blocks” were then searched for proxies using an r^2^ threshold of 0.8 and a distance limit of 250 kilobases, resulting in “r^2^ > 0.8 LD blocks” defining potentially causal variants at each locus. The “r^2^ > 0.8 LD blocks” for the ALT results were then compared to the “r^2^ > 0.8 LD blocks” for the AST results, and any cases where these blocks shared at least one variant between the two GWAS were treated as potentially colocalized association signals between the two GWAS. In these cases, a representative index variant was chosen to represent the results of both GWAS by choosing the index variant of the GWAS with the more significant p-value. Next, these putative colocalized association signals were then distance pruned by iteratively removing neighboring index variants within 500 kilobases of each index variant with less significant p-values (the minimum p-value between the two GWAS was used for the distance pruning procedure.) The Manhattan plot of METAL results with labeled colocalization signals was created using the CMplot R package^113^.

### Annotation of associated loci and variants

Index variants and their corresponding strongly-linked (r^2^ > 0.8) variants were annotated using the following resources: distance to closest protein-coding genes as defined by ENSEMBL v98 using the BEDTools package^114^, impact on protein-coding genes using the ENSEMBL Variant Effect Predictor (VEP) software package^115^ with the LOFTEE plugin to additionally predict protein-truncating variants^116^; eQTLs (only the most significant eQTL per gene-tissue combination) from GTEx v8 (obtained from the GTEx Portal) for liver, kidney cortex, and skeletal muscle^117^; a published meta-analysis of four liver eQTL studies^41^; the eQTLGen meta-analysis of blood eQTL studies^118^; and GWAS results from the NHGRI-EBI GWAS Catalog (r2020-01-27)^119^, filtered to associations with p < 5 × 10^−8^.

### Association of ALT- and AST-associated loci with liver disease

Index variants were tested for association with any liver disease using ICD10 codes K70-K77 in inpatient hospital diagnoses, causes of death, and GP clinical event records, using SAIGE, with the same covariates used for the liver enzymes (age, sex, and genetic PCs 1-12) plus a covariate for each of the following: whether the subject was recruited in Scotland, whether the subject was recruited in Wales, and whether the patient had GP clinical event records available. Association results were meta-analyzed across the four sub-populations using METAL using the default method (combining p-values) to obtain the final p-value. To obtain effect sizes and standard errors, the same procedure was performed but using PLINK (on the unrelated subset of each population) and using the classical method in METAL (combining effects and standard errors.)

### Sequencing-based validation of rs188273166 array genotyping

Whole exome sequencing was available for 301,473 of the 487,327 array-genotyped samples. DNA was extracted from whole blood and was prepared and sequenced by the Regeneron Genetics Center (RGC). A complete protocol has been described elsewhere^120^. Briefly, the xGen exome capture was used and reads were sequenced using the Illumina NovaSeq 6000 platform. Reads were aligned to the GRCh38 reference genome using BWA-mem^121^. Duplicate reads were identified and excluded using the Picard MarkDuplicates tool (Broad Institute)^122^. Variant calling of SNVs and indels was done using the WeCall variant caller (Genomics Plc.)^123^ to produce a GVCF file for each subject (GVCF files are files in the VCF Variant Call Format that are indexed for fast processing). GVCF files were combined to using the GLnexus joint calling tool^124^. Post-variant calling filtering was applied as described previously^120^.

### Replication of *SLC30A10* Thr95Ile associations

ALT and AST association tests were repeated as described for the genome-wide scans, using SAIGE and PLINK, in the “Other White” and “White Irish” populations, for the *SLC30A10* Thr95Ile (rs188273166) variant. In the two DiscovEHR Geisinger Health Service (GHS) cohorts, association tests were performed using BOLT^125^ with covariates for age, age squared, age x sex, sex, and the first ten principal components of genetic ancestry; ALT, AST, and hematocrit values were taken from the median of lab values available. Results were meta-analyzed across the four populations. A forest plot was created using the forestplot package in R^126^.

### Testing linkage of *SLC30A10* Thr95Ile to common GWAS variants

To test linkage of *SLC30A10* Thr95Ile (rs188273166) to common GWAS variants, the GWAS Catalog was searched for all results where “Mapped Gene” was assigned to *SLC30A10*; because of the very relevant phenotype, blood Mn-associated variant rs1776029, an association that is not in the GWAS Catalog, was also included in the analysis, as well at cT1-associated variant rs759359281. LD calculations were performed in PLINK, using the White British unrelated subpopulation, between rs188273166 and the GWAS variants with the options --r2 dprime-signed in-phase with-freqs --ld-window 1000000 --ld-window-r2 0. For rs1776029, an additional Fisher’s exact test was performed to determine the confidence interval of the enrichment of rs188273166 on the rs1776029 haplotype. The linked alleles from PLINK were then used in conjunction with the effect allele from the reported papers to determine the direction of effect. The GWAS Atlas website^46^ was used (the PheWAS tool) to determine the direction of effect for the linked alleles from the original paper; in cases where the original paper from the GWAS Catalog did not report a direction of effect, other papers for the same phenotype and variant from GWAS Atlas were used to determine the direction of effect and cited accordingly (**Supplementary Table 13**). Reference epigenome information for the GWAS variants was obtained by searching for rs1776029 in HaploReg v4.1^127^.

### Phenome-wide association study of *SLC30A10* Thr95Ile

A phenome-wide association study of *SLC30A10* Thr95Ile (rs188273166) was performed by running SAIGE and PLINK against a set of ICD10 diagnoses and quantitative traits, obtained as described above, and using the covariates described above for the test of association with liver disease. ICD10 diagnoses were filtered to include only those at a three-character (category), four-character (category plus one additional numeral), or “block” level that were frequent enough to test in both subpopulations and without significant collinearity with the sex, GP availability, or country of recruitment covariates: at least 100 diagnoses overall, and at least one diagnosis in each of the following subgroups, to avoid collinearity with covariates while running SAIGE: the with- and without-GP data subgroups, men, women, and each of the three recruitment countries. This resulted in 4,397 ICD10 codes to test, serving as the multiple hypothesis burden.

### Bioinformatic analysis of *SLC30A10* Thr95Ile

To visualize Thr95Ile on the protein sequence of SLC30A10, UNIPROT entry Q6XR72 (ZN10_HUMAN) was accessed^72^. In UNIPROT, natural variants causing HMNDYT1^32,34,56^ and mutagenesis results^56,73,128^ were collated from the literature and highlighted. CADD score v1.5^129^ was downloaded from the authors’ website. SIFT score was obtained from the authors’ website using the “dbSNP rsIDs (SIFT4G predictions)” tool^130^. PolyPhen score and multiple species alignment was obtained from the authors’ website using the PolyPhen-2 tool^131^.

### Immunofluorescence of SLC30A10 localization in cultured cells

HeLa cells (ATCC®, Manassas, VA) were grown in Eagle’s Minimum Essential Medium (ATCC®, Manassas, VA) containing 10% fetal bovine serum (Gibco, Carlsbad, CA) at 37°C and 5% CO_2_. All plasmid transfections were performed using Lipofectamine™ 2000 (Invitrogen, Carlsbad, CA) and Opti-MEM (Gibco, Grand Island, NY) according to manufacturer’s specifications. FLAG-tagged SLC30A10 plasmid constructs designed with a linker sequence in pCMV6-AN-3DDK (Blue Heron Biotech, Bothell, WA) included wild type, del105-107, L89P, T95I, and used an empty vector for one of the negative controls.

HeLa cells were grown on 8-chambered slides for 48 hours post-transfection. IF procedures were performed at room temperature unless otherwise noted. HeLa cells were rinsed in dPBS (Gibco, Grand Island, NY), fixed with 4% paraformaldehyde (in water) (Electron Microscopy Sciences, Hatfield, PA) for 10 min, rinsed in 4°C PBS (Invitrogen, Vilnius, LT), and permeabilized for 5 minutes with 0.1% Triton X-100 (Sigma-Aldrich, St. Louis, MO). After rinsing in PBS and blocking in 2% BSA (in PBS) (Jackson ImmunoResearch, West Grove, PA) for 30 minutes, the cells were stained with 2% BSA blocking solution containing monoclonal ANTI-FLAG® M2-FITC, Clone M2 (dilution 1:100; Sigma-Aldrich, St. Louis, MO) and Calnexin Monoclonal Antibody, Clone AF18 (dilution 1:100; Invitrogen, Carlsbad, CA). After three final washes in dPBS, mounting medium with DAPI (Vector Laboratories, Burlingame, CA) was added and sealed under a coverslip with nail polish. Images were captured with the REVOLVE Echo microscope at 20X magnification.

## Acknowledgements

This research has been conducted using the UK Biobank resource, application number 26041. We thank the UK Biobank participants for their donations to this resource.

## Author Contributions

A.D., A.F.C., S.T., L.W., M.P., and P.N. performed the main computational analyses; C.Q. and H.C.T. performed experiments; L.L., N.V., and M.F. performed replication analyses in DiscovEHR; L.W. wrote the manuscript; all authors interpreted results and edited the manuscript.

## Ethical Compliance

Ethics oversight for the UK Biobank is provided by an Ethics and Governance Council which obtained informed consent from all participants for health-related research. All research described was performed within the framework of Application 26041.

## Data Availability

Complete summary statistics from the genome-wide association studies of ALT, AST, and extrahepatic bile duct cancer have been submitted to the NHGRI-EBI GWAS catalog (https://www.ebi.ac.uk/gwas/).

## Supplementary Figures

**Supplementary Figure 1:**
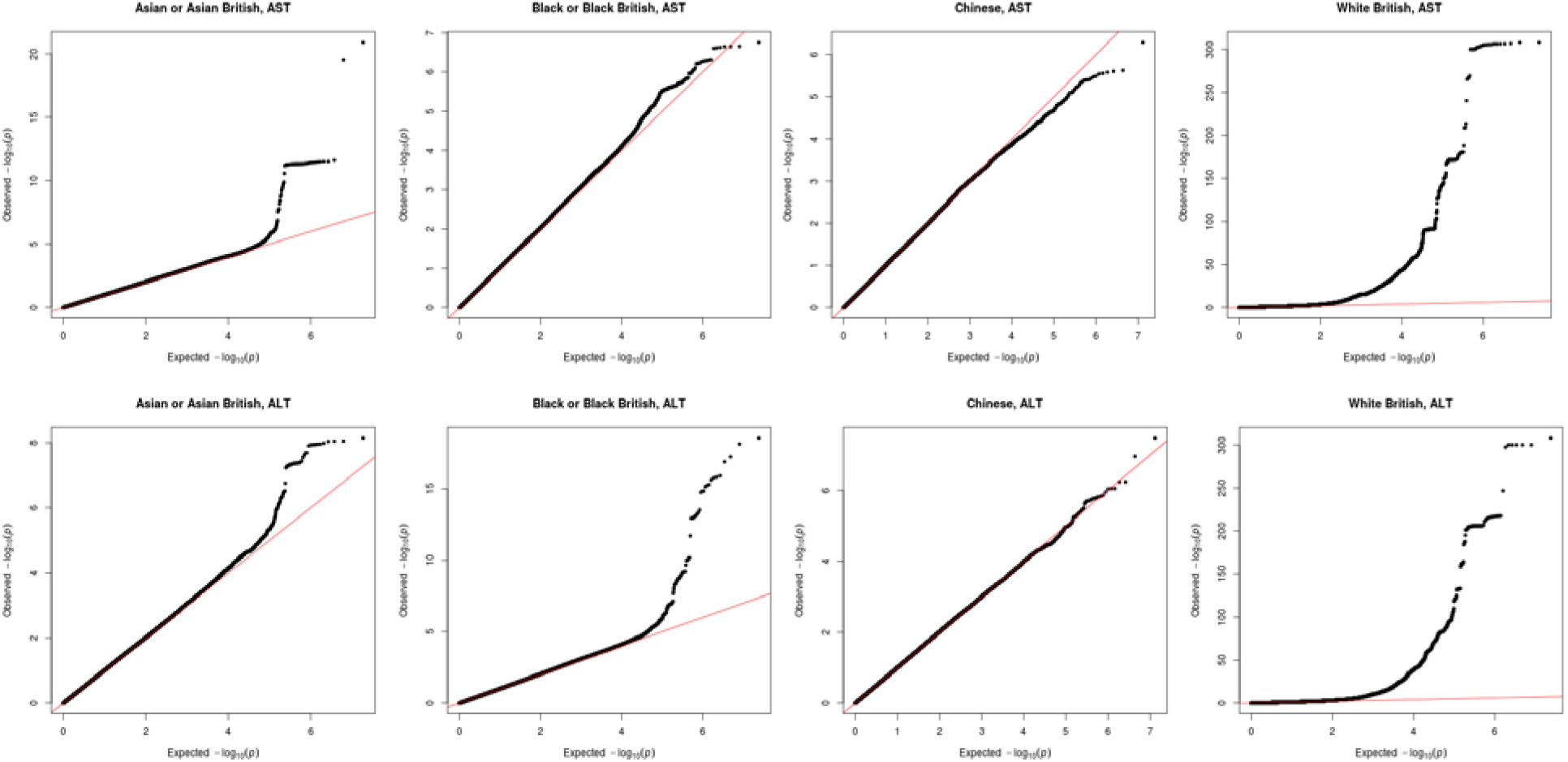
Q-Q plots of GWAS p values (from SAIGE) for each sub-population and enzyme.

**Supplementary Figure 2:**
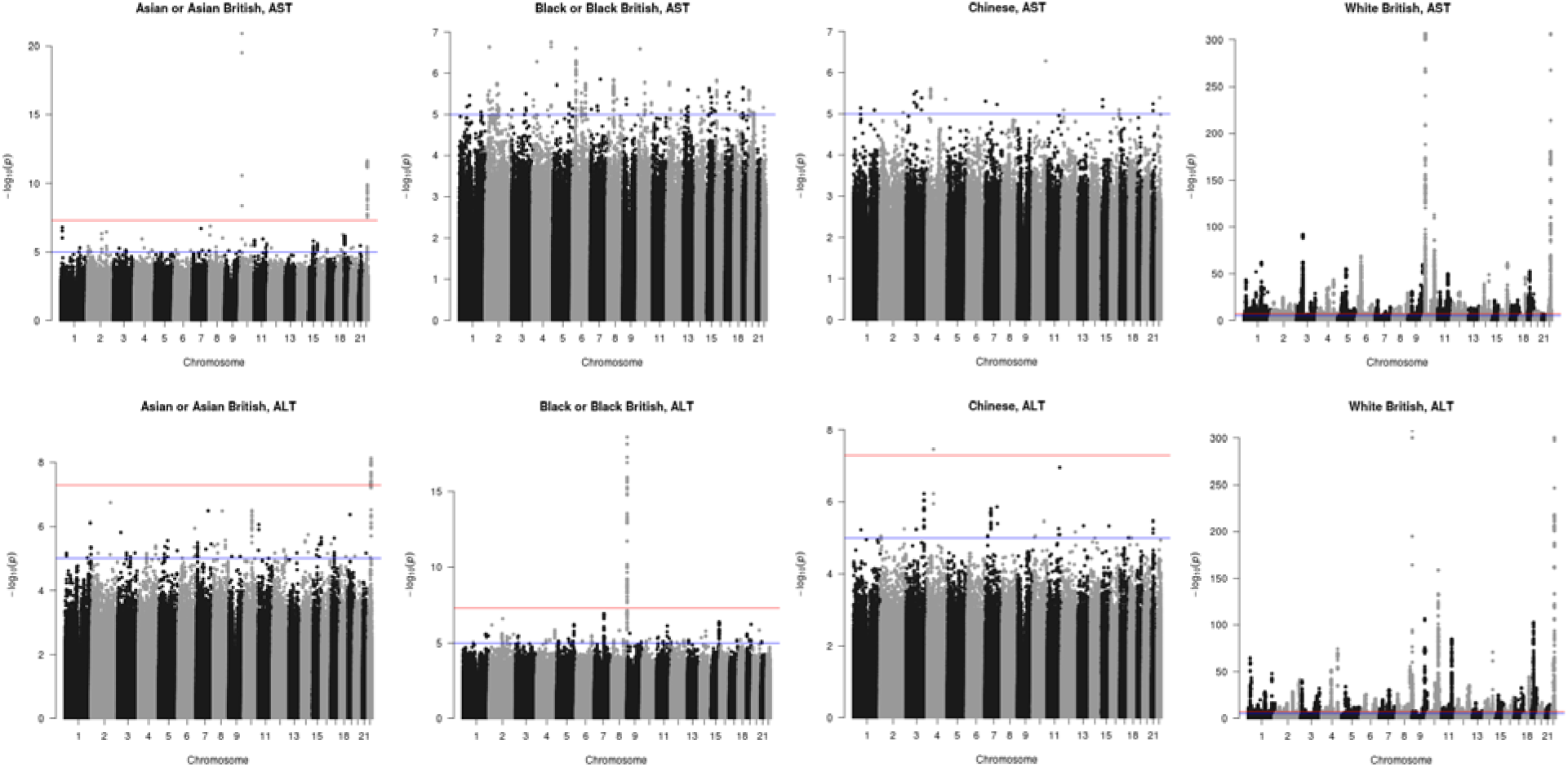
Manhattan plots of GWAS p values (from SAIGE) for each sub-population and enzyme.

**Supplementary Figure 3:**
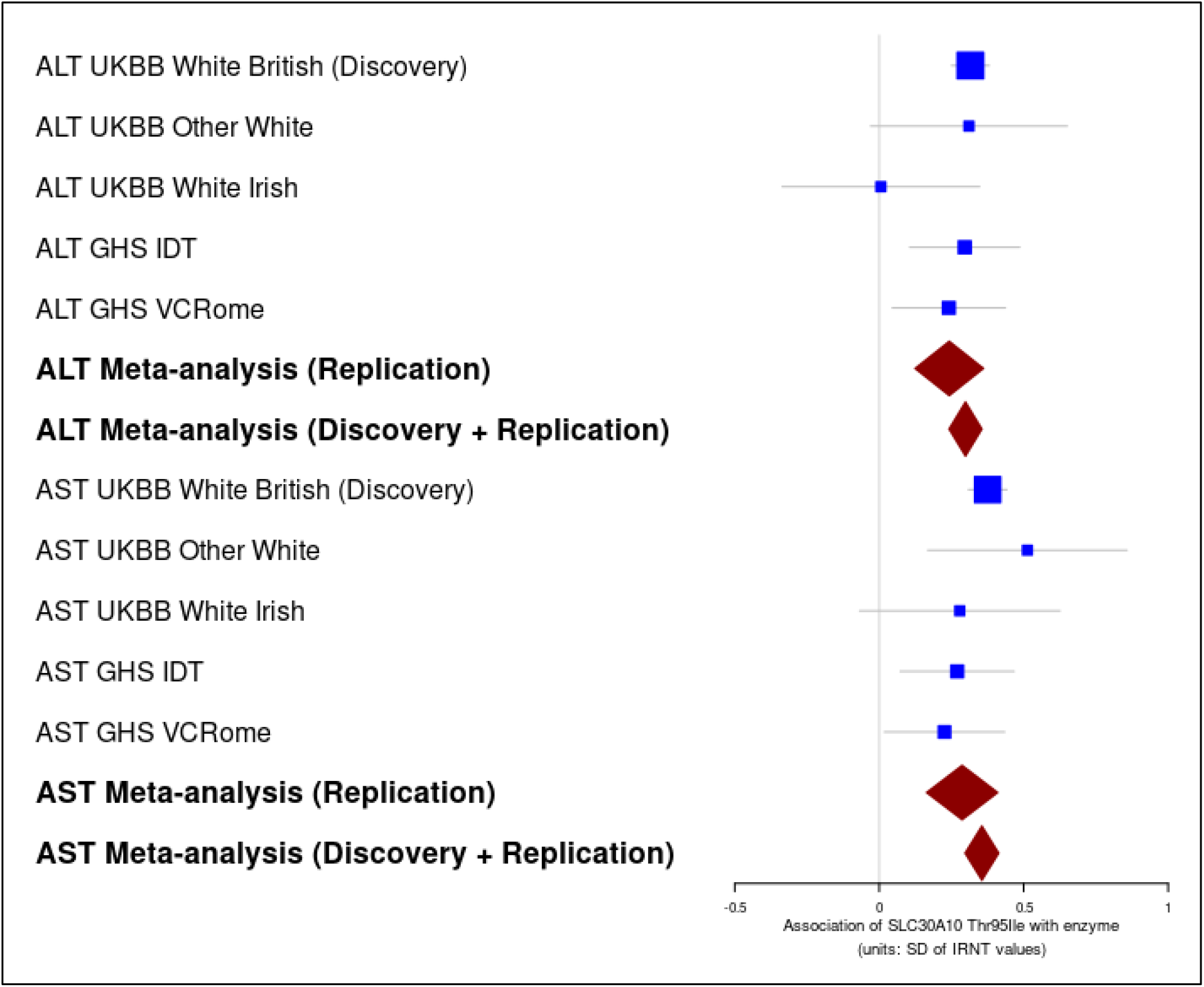
Forest plot of *SLC30A10* Thr95Ile (rs188273166) association with ALT and AST in the discovery and replication groups. Boxes show effect size estimates from PLINK, and error bars show 95% confidence intervals.

**Supplementary Figure 4:**
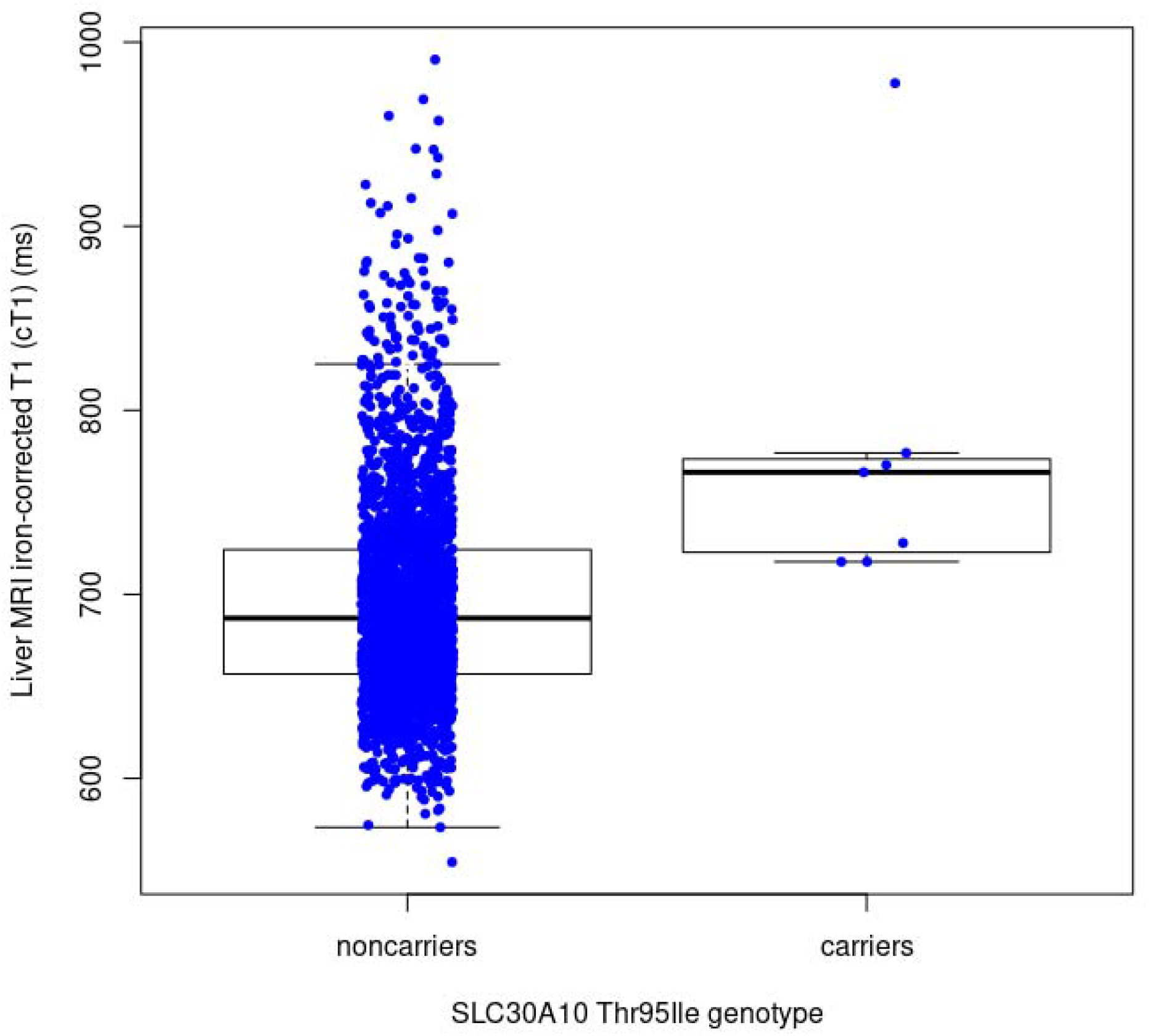
iron-corrected T1 (cT1) values from liver MRI in seven SLC30A10 Thr95Ile carriers (limited to the White British population).

**Supplementary Figure 5:**
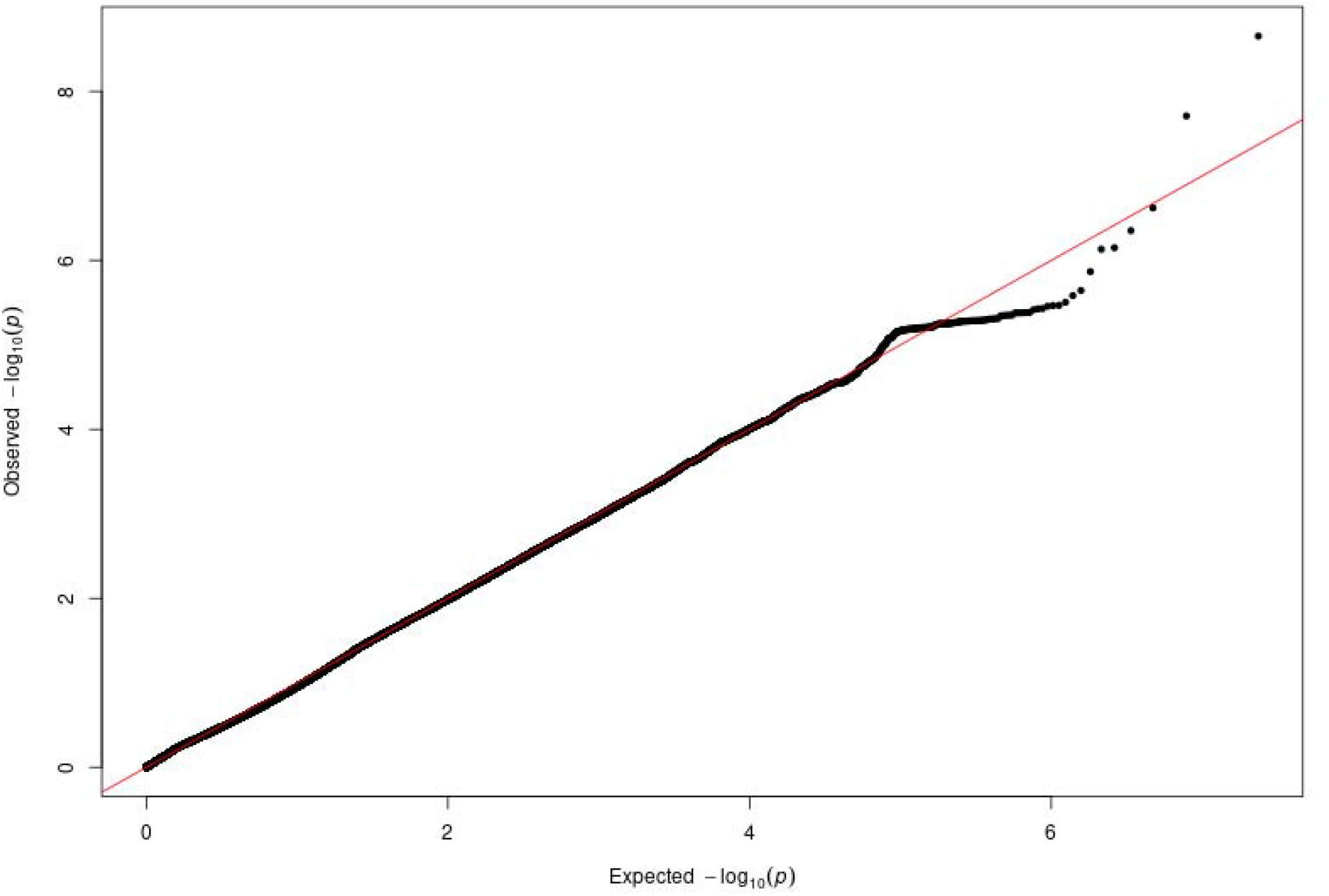
Q-Q plot of GWAS (p value from SAIGE) for ICD10 diagnosis C24.0, extrahepatic bile duct cancer.

**Supplementary Figure 6:**
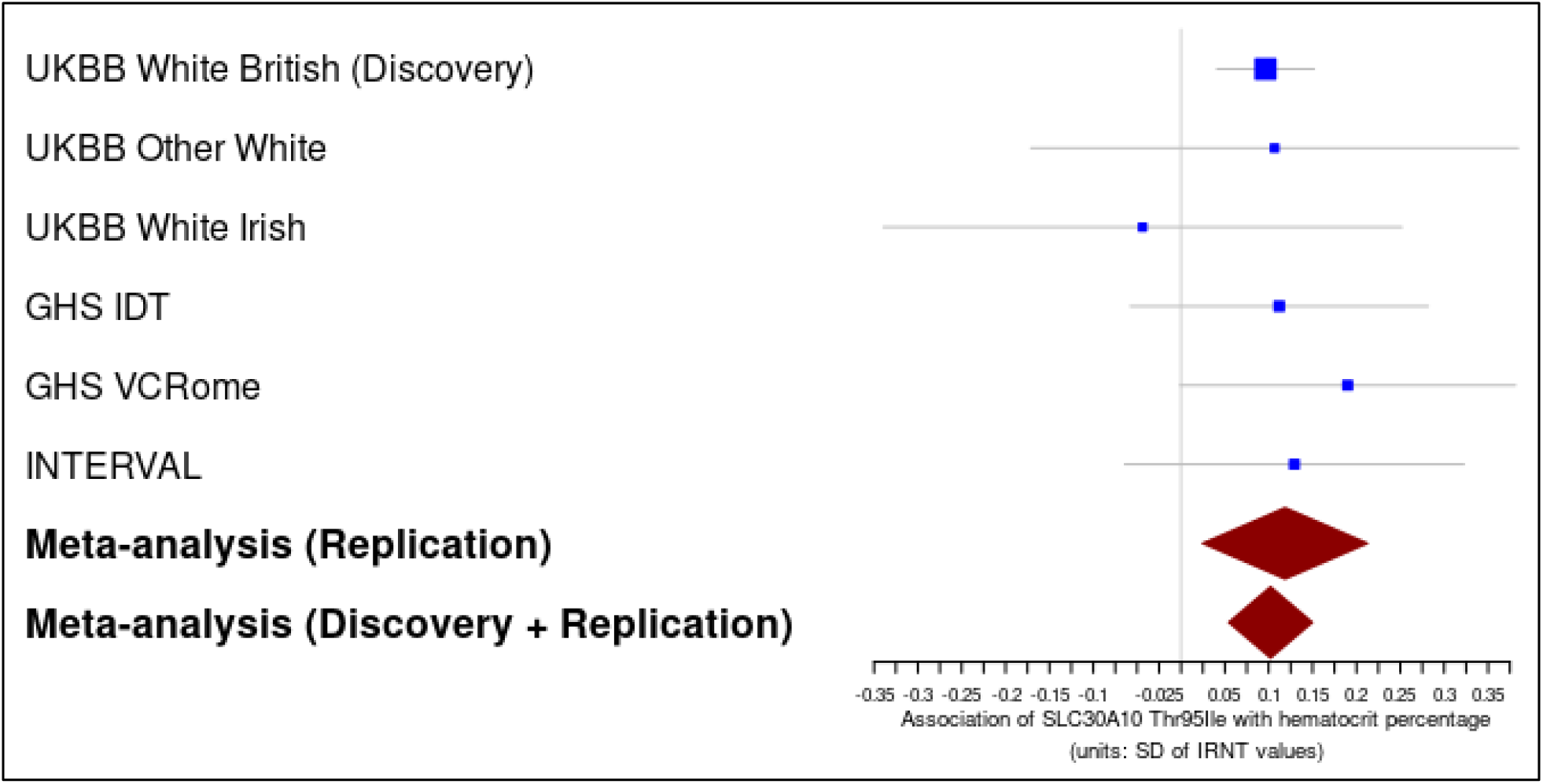
Forest plot of *SLC30A10* Thr95Ile (rs188273166) association with ALT and AST in the discovery and replication groups. Boxes show effect size estimates from PLINK, and error bars show 95% confidence intervals.

**Supplementary Figure 7:**
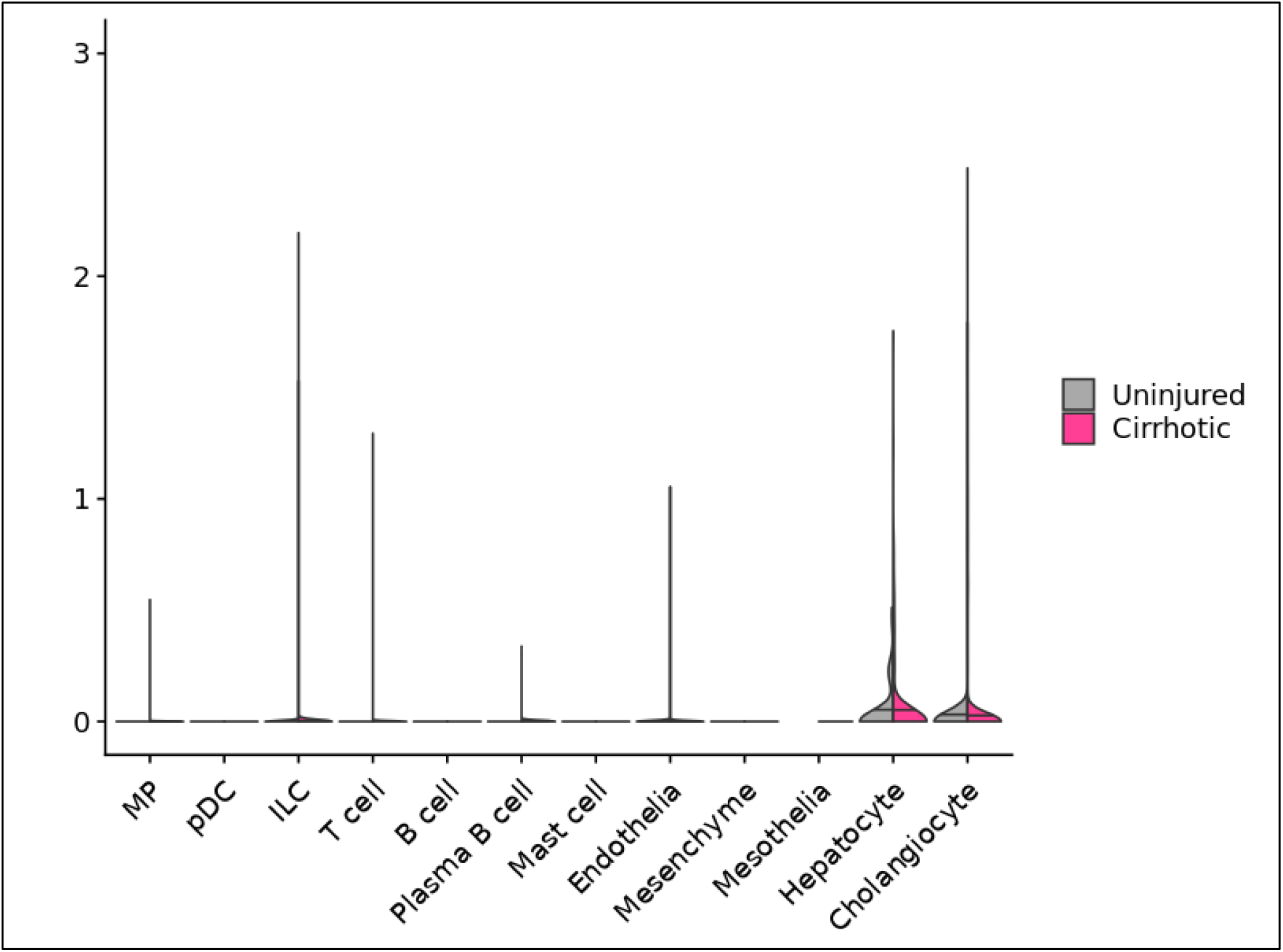
Single-cell expression data, showing distribution of expression values for SLC30A10 from Ramachandran et al.^53^ Produced by https://www.livercellatlas.mvm.ed.ac.uk/. *

**Supplementary Figure 8:**
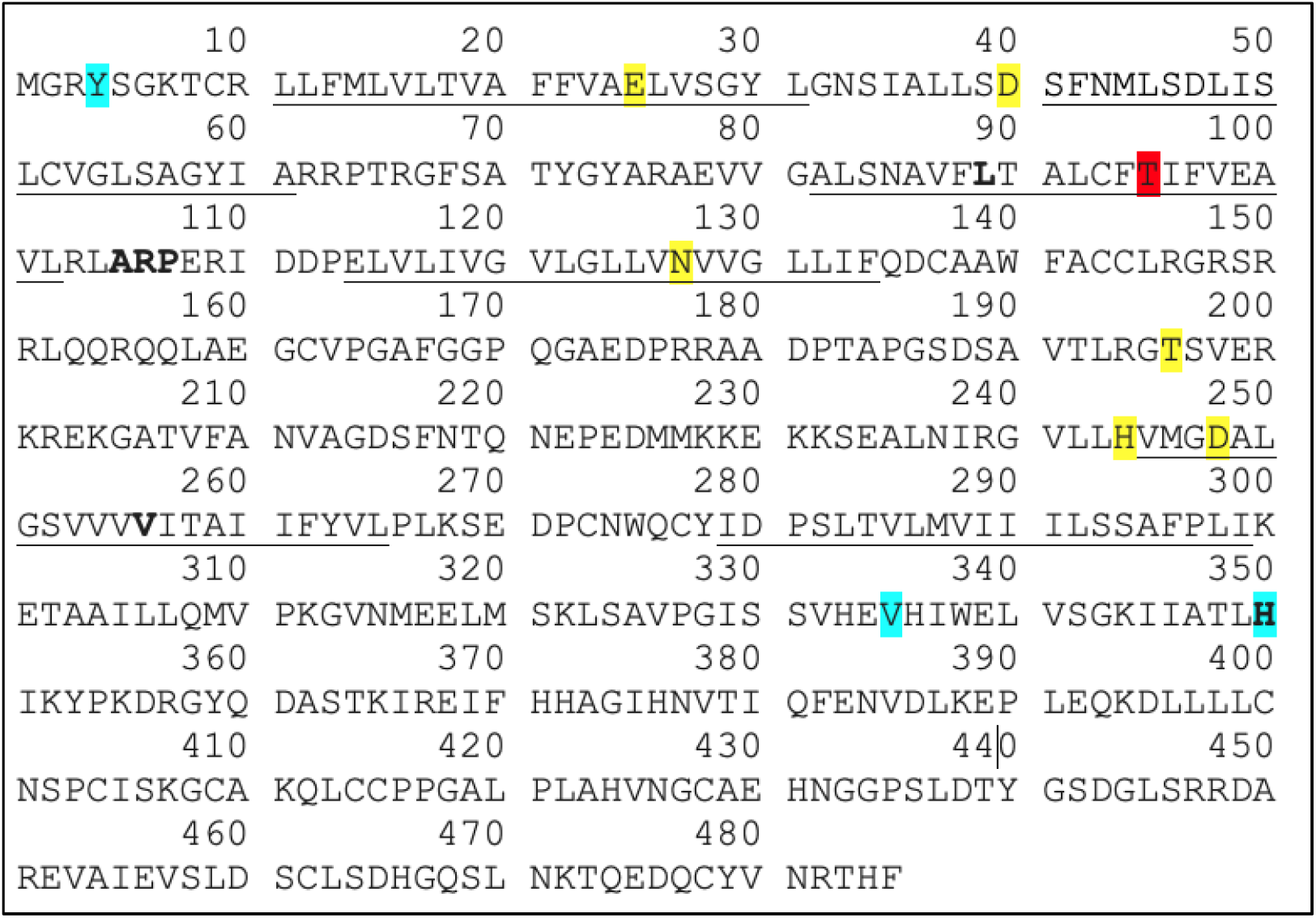
Protein sequence of SLC30A10 along with experimental evidence cited by UNIPROT^72^. Underlined are the six transmembrane domains. Highlighted red is Thr95Ile. Highlighted in yellow are variants that demonstrated abolished Mn transport or membrane localization in vitro; highlighted in blue are variants that demonstrated lesser or no effect on Mn transport or membrane localization in vitro^56,73,128^. In bold are variants known to cause SLC30A10 deficiency (HMNDYT1)^32,34,56^.

